# Schema formation in a neural population subspace underlies learning-to-learn in flexible sensorimotor problem-solving

**DOI:** 10.1101/2021.09.02.455707

**Authors:** Vishwa Goudar, Barbara Peysakhovich, David J. Freedman, Elizabeth A. Buffalo, Xiao-Jing Wang

## Abstract

Learning-to-learn, a progressive speedup of learning while solving a series of similar problems, represents a core process of knowledge acquisition that draws attention in both neuroscience and artificial intelligence. To investigate its underlying brain mechanism, we trained a recurrent neural network model on arbitrary sensorimotor mappings known to depend on the prefrontal cortex. The network displayed an exponential time course of accelerated learning. The neural substrate of a schema emerges within a low-dimensional subspace of population activity; its reuse in new problems facilitates learning by limiting connection weight changes. Our work highlights the weight-driven modifications of the vector field, which determines the population trajectory of a recurrent network and behavior. Such plasticity is especially important for preserving and reusing the learnt schema in spite of undesirable changes of the vector field due to the transition to learning a new problem; the accumulated changes across problems account for the learning-to-learn dynamics.

## Introduction

In Psychology, *Schema* refers to an abstract mental representation that we deploy to interpret and respond to new experiences, and to recall these experiences later from memory [1, 2]. Mental schemas are thought to express knowledge garnered from past experiences [2, 3, 4]. For example, expert physicists apply relevant physics schemas when they categorize mechanics problems based on governing physical principles (e.g. conservation of energy or Newton’s second law); by contrast, novice physicists who lack these schemas resort to categories based on concrete problem cues (e.g. objects in the problem or their physical configuration) [5]. What is the brain mechanism of schemas, and what makes it essential for rapid learning and abstraction?

One type of schema is called a “learning set”. In a pioneering experiment, H. F. Harlow trained macaque monkeys on a series of stimulus-reward association problems [6]. While keeping the task structure fixed, each problem consisted of two novel stimuli that had to be correctly mapped onto rewarded versus nonrewarded, respectively. Harlow found that the monkeys progressively improved their learning efficiency over the course of a few hundred problems, until they could learn new problems in one shot. He concluded that rather than learning each problem independently of the earlier ones, the monkeys formed an abstract learning set that they deployed to learn new problems more efficiently — they were *learning-to-learn*. Evidence of schema formation from prior knowledge has been demonstrated in humans [7, 8] and nonhuman animals [9, 10, 11]. Moreover, converging lines of evidence derived from functional connectivity [12, 13], structural plasticity [13], lesion [9], pharmacological blockade [14], and gene expression [15] studies, attribute the acceleration of learning to the rapid integration of new experiences into pre-existing schema that are encoded in the neocortex. This has motivated a neurocentric definition of a schema as a network of strongly interconnected neocortical representations that affect processing of new information [7, 12, 16]. However, these previous experiments did not elucidate *how*, mechanistically, a neural circuit realizes a schema and expedites learning.

Schemas are posited to emerge as an abstraction of the commonalities across previous experiences [4, 17]. It is the generalization of these abstract representations to novel situations that is believed to accelerate learning [18, 19, 20]. Indeed, the abstract neural representation of shared task variables has been observed across consecutively learned problems when experience on earlier problems facilitates later learning [21, 22]. Furthermore, the progressive improvement in learning efficiency observed by Harlow suggests that this process of abstract representation-facilitated learning undergoes progressive refinement. The structure learning hypothesis [23] equates learning to a change in the brain’s internal parameters which control behavior, and posits that the progressive improvement in learning efficiency emerges with a low-dimensional task-appropriate realization of the internal parameter space. Parameter exploration within such a space is less demanding, which makes learning more efficient. Therefore, while schema formation emphasizes an abstraction of the task’s structure, structure learning emphasizes learning how to efficiently use a schema to aid in generalization.

In spite of tremendous progress in machine intelligence, learning-to-learn presents a major challenge in presently available artificial systems. Machine learning studies have proposed *meta-learning* approaches wherein model parameters that promote rapid generalization to new problems are explicitly favored and sought [24, 25]. Yet, it is not known whether such mechanisms are necessary computationally or present in the brain. Can learning-to-learn arise solely from the natural dynamics of learning?

We explored this question of broad interest to brain research, cognitive science and artificial intelligence, by examining the neural mechanisms of learning-to-learn in recurrent neural network (RNNs). As a behavioral paradigm we chose learning of arbitrary sensorimotor associations, which requires the learning of arbitrary mappings between sensory stimuli and motor consequents [26, 27], and is essential for flexible behavior [28]. Macaque monkeys exhibit learning-to-learn on association problems, they learn new problems within an average of 20 trials when they are well-trained [29]. Furthermore, their prefrontal cortex is causally engaged during rapid problem learning. Prefrontal neurons represent task stimuli and responses during visuomotor association trials [26, 29]. Prefrontal lesions produce substantial visuomotor association learning deficits [28, 30, 31]. We sought to understand whether and how a sensorimotor association schema may be encoded by these prefrontal representations, how it is applied to new problems, and how its usage is refined to improve learning efficiency.

Towards this end, we trained and analyzed an RNN model of prefrontal computation and learning. We found that RNNs trained on a series of sensorimotor association problems exhibit robust learning-to-learn despite the absence of meta-learning: the number of trials to learn a problem decays exponentially with the number of previously learned problems without an explicit mechanism to accelerate learning with increasing experience. We analyzed the population activity of the RNN’s units via subspace decomposition to uncover population-level latent variable representations [10, 32], and we used manifold perturbations to study the causal relationship between learning efficiency and the reuse of existing population representations to learn [33]. The analyses revealed that the model develops neural correlates of the task’s schema — a low-dimensional neural manifold that represents shared task variables in an abstract form across problems. Its reuse avoids the formation of representations *de novo* while learning problems which considerably accelerates learning by limiting the connection weight changes required to learn them. We introduce a novel measure relating these weight modifications to population activity changes, which we term the *weight-driven vector field change*. This measure showed that the reused representations are not entirely invariant across problems. Instead, mapping new stimuli can modify the reused representations in undesirable ways. Connection weight changes are primarily recruited to prevent such modifications. Moreover, the weight changes in early problems improve the invariance of the reused representations, limiting the degree to which they would be modified in the future, which further accelerates learning. The accumulation of such improvements over a series of problems supports structure learning and promotes learning-to-learn.

## Results

### Trained neural network models exhibit learning-to-learn without meta-learning

We evaluated whether an RNN model could demonstrate learning-to-learn of delayed sensorimotor association problems. In each problem, the model learned a unique mapping between a pair of sensory stimuli (e.g. images) and a pair of motor responses (Fig. 1a). Each trial began with a 0.5 second sample epoch, when a sensory stimulus was presented together with a fixation cue, and the model was required to maintain fixation. A 1 second delay epoch followed, when the model had to continue fixation in the absence of the sample stimulus. The trial concluded with a 0.5 second choice epoch signalled by removal of the fixation cue, when the model had to report its choice of an appropriate motor response. The two sample stimuli in each problem were randomly generated. The model was composed of a population of recurrently (or laterally) connected firing rate units that received eleven inputs, one signalling fixation and ten signalling features of a sample stimulus (Fig. 1b). Such stimulus representations are consistent with a recent finding that visual objects are represented in the monkey inferotemporal cortex by a feature-based topographic map [34]. The model is also consistent with lesion studies which demonstrate the causal involvement of inferotemporal-prefrontal connections in visuomotor learning and retention [30, 35]. Response choices were read out from the population’s activity by three output units that represented fixation, motor response 1, or motor response 2.

**Figure 1:**
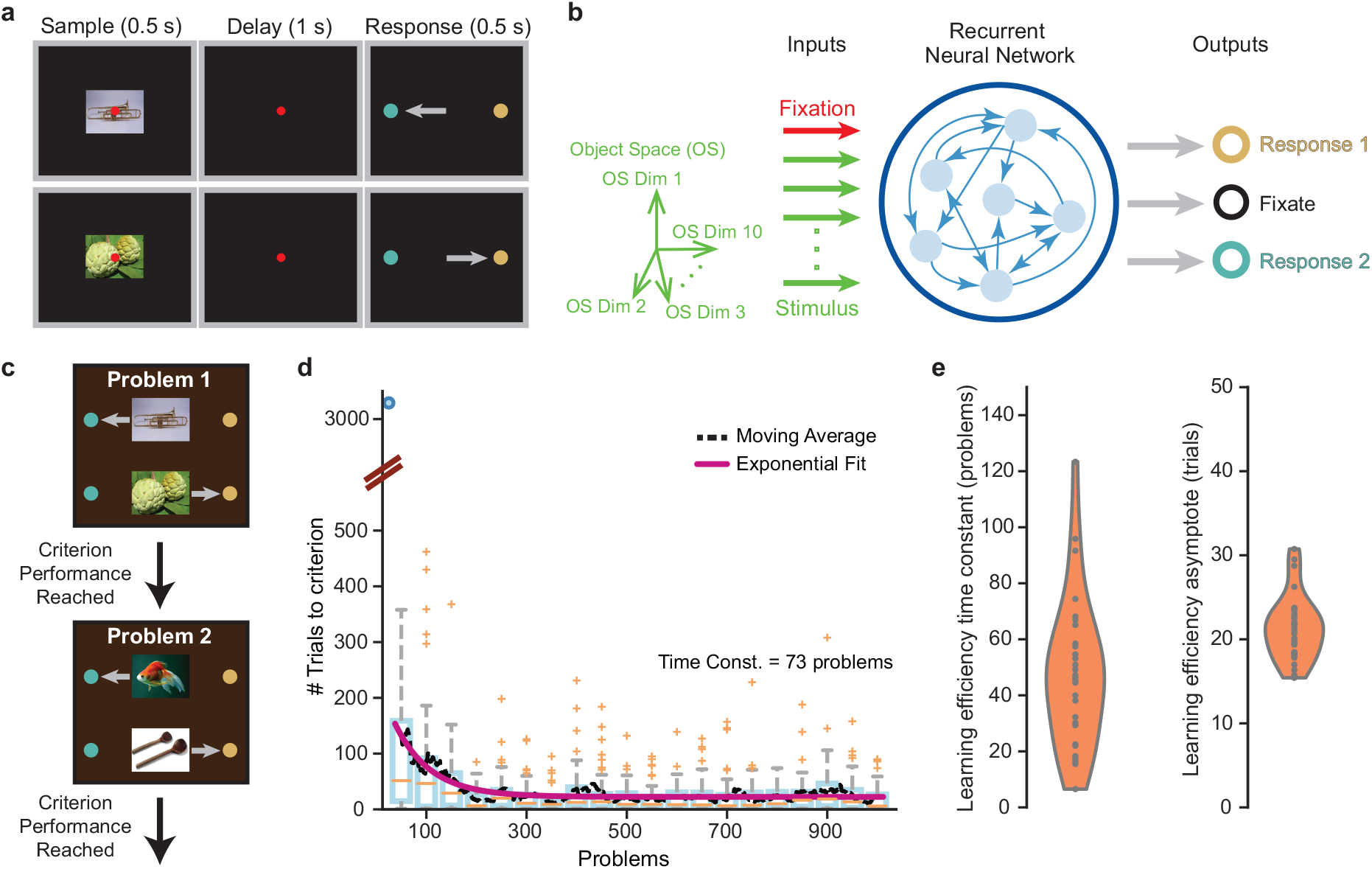
Recurrent neural networks trained on a series of delayed sensorimotor association problems exhibit learning-to-learn. **a.** An example problem illustrating the structure of the delayed sensorimotor association task. The model must learn to associate each of two sensory stimuli (e.g. images) with a corresponding motor response (e.g. a saccade). Targets are colored to emphasize the distinction between response choices, and not to indicate that the response targets are colored. **b.** RNN model trained to perform the task. It is composed of recurrently connected rate units that receive a fixation stimulus and features of the sample sensory stimulus as inputs, and reports its response choices via output units corresponding to fixation, motor response choice 1 (brown), or motor response choice 2 (teal). **c.** Learning-to-Learn training protocol. The model is trained on a series of sensorimotor association problems, each with randomly chosen sample stimulus pair. It is transitioned to a new problem once it reaches criterion performance on the current problem. **d.** Learning efficiency, measured as the number of trials to criterion performance on a problem, over a series of 1000 problems learned by a network. Box plots summarize the learning efficiency in groups of 50 consecutive problems. The number of trials to criterion on a problem decreases with the number of previously learned problems. This is characterized by a decaying exponential function and demonstrates the model’s ability to produce learning-to-learn. **e.** 30 RNNs with different initial conditions exhibit robust learning-to-learn, as indicated by the time-constants (left) and asymptotes (right) of the exponential fits to their learning efficiency over a series of 1000 problems.

The model was trained on a problem one trial at a time. Its parameters were adjusted at the end of each trial to minimize the errors in its output responses, until the output responses achieved criterion accuracy (see *Methods*). The model was then transitioned to a new problem (Fig. 1c). Crucially, training was performed without an explicit meta-learning objective. A network trained on a series of delayed sensorimotor association problems demonstrated learning-to-learn (Fig. 1d). The network required a few thousand trials to learn the first problem, which was expected because the network was initialized with random connection weights. By contrast, solving the second problem took only a few hundred trials. Thereafter the trials-to-criterion progressively decreased over the next few hundred problems, plateauing at an average of 20 trials per problem. This decrease was well-fit by a decaying exponential function, which closely matched a 30-problem moving average of the network’s trials-to-criterion. This performance is commensurate with learning-to-learn in macaque monkeys, which exhibit an exponential decrease in their trials-to-criterion when trained on a series of association problems [36], and demonstrate learning within 15-20 trials when well-trained [29]. The fit’s parameters quantify the network’s learning-to-learn performance: the time constant measures how quickly it produces learning-to-learn, and the learning efficiency asymptote measures its trials-to-criterion plateau. We note that while naive monkeys undergo behavioral shaping on the desired response set before they are introduced to the task, a naive network’s learning efficiency on the first problem reflects learning both to generate basic responses, and the specifics of the problem. To avoid this confound related to learning the response set, we quantified the network’s learning-to-learn performance starting with the second problem.

We tested the robustness of these results by similarly training 30 independently initialized networks. Across these networks, the learning-to-learn time constants and asymptotes were limited to a narrow range (Fig. 1e; time constant mean=47.52, std. dev.=26.22; asymptote mean=21.33, std. dev=3.85). We further tested the model over a range of hyper-parameter settings (f-I transfer functions, learning rates, weight and firing rate regularization levels), and observed robust learning-to-learn across all conditions (Supplementary Fig. 1). In addition, we found that the model was faster at re-learning problems after subsequently learning several new problems (Supplementary Fig. 2), suggesting that it retains a memory of previously learned problems. Taken together, these results demonstrate that networks trained on a series of delayed sensorimotor association problems robustly exhibit learning-to-learn, despite the absence of an explicit meta-learning objective.

### Abstracted neural manifold governs the task’s schema and drives output responses

The activity of a recurrently connected population of *N* units co-evolves over the duration of a trial, forming a trajectory in an *N*-dimensional population state space (Fig. 2a, top). When a problem is learned, the network responds to each sample stimulus with a trajectory that appropriately subserves stimulus integration, decision making, working memory maintenance, and fixation/motor response choice. We demixed [37] (see *Methods*) population trajectories from consecutively learned problems to identify shared representations, if any, of the latent variables that support these computations. This procedure decomposed the trajectories into components embedded within two non-overlapping subspaces of the population state space (Fig. 2a, middle): Decision representations embedded within the *decision subspace* revealed similarities between trajectories that shared target responses; stimulus representations embedded within the *stimulus subspace* varied in a problem- and sample stimulus-dependent manner. We further decomposed the two decision representations in each problem into a mean decision representation, where the mean was taken over both decision representations (Fig. 2a, bottom left), and residual decision representations given by subtracting out this mean from the two decision representations (Fig. 2a, bottom right).

**Figure 2:**
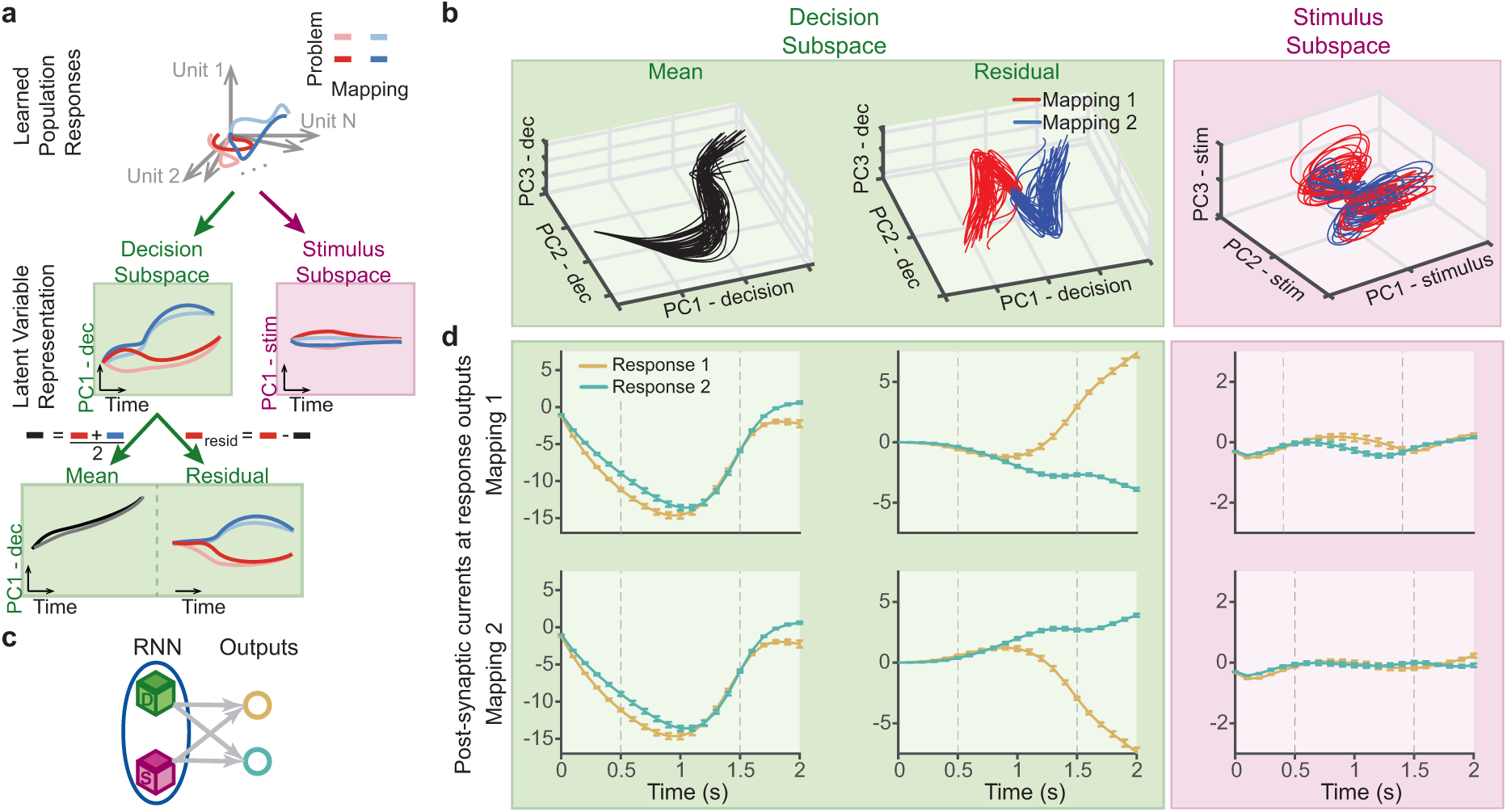
Neural representations of decision and choice are shared across problems. **a.** Schematic of the demixing procedure that identifies shared versus problem-dependent components of the neural representations. Population trajectories for the two mappings in 50 consecutively learned problems (illustrated for 2 problems, for clarity) are decomposed into components within a decision subspace which are shared by trajectories that map their respective sample stimuli onto a common response choice, and problem-dependent components embedded in a stimulus subspace. The shared decision representations are further decomposed into their mean and residual components for each problem. **b.** Decomposed representations for problems 1-50, presented in the first 3 principal components of their respective subspaces. **c.** Schematic illustrating that the component representations collectively drive the response choice outputs. **d.** The net current from the mean (left) and residual (center) decision representations, and the stimulus representations (right), to response 1 (brown) and response 2 (teal) outputs, in mapping 1 (top) and mapping 2 (bottom) trials. The mean decision components inhibit motor responses during the sample and delay epochs, and the residual decision components drive the correct response while inhibiting the incorrect one. Dashed vertical lines indicate the end of the sample and delay epochs. Plots show mean of the net currents across the 50 problems, and error bars indicate their standard errors.

Decomposing the trajectories from the first 50 consecutively learned problems in this manner revealed a low-dimensional shared decision subspace (mean = 2.36 dimensions; std. dev. = 0.18 dimensions across 10 networks), whose constituent decision representations explained most of the variance in population activity across problems (mean = 88.54%; std. dev. = 3.16% across 10 networks). Furthermore, the mean decision representations lay close to each other in state space, forming a shared manifold across problems (Fig. 2b, left). The residual decision representations consistently encoded the decision and choice of either response across problems, thus forming a shared manifold for each decision (Fig. 2b, center). The persistence of a low-dimensional shared manifold which explains a majority of the population’s variance across problems demonstrates a strong abstraction of the shared task variables that it encodes. It bears mentioning that the model retains and reuses this manifold across problems, despite changes in the stimulus set and the weight change-induced change in network dynamics that transpires while learning. Moreover, population activity changes during learning are largely determined by changes in these shared representations (Supplementary Fig. 3). In contrast, the stimulus representations (Fig. 2b, right) were higher dimensional (mean = 7.98 dimensions; std. dev. = 1.48 dimensions across 10 networks), but explained a small proportion of the population variance. Interestingly, the distribution of neural activity in population state space at the beginning and end of problem learning closely resemble each other (Supplementary Fig. 4). These results demonstrate that the model even reuses pre-established representations when responding to novel sample stimuli and learning their mappings.

Next, we examined the relative contribution of these components to the output responses by measuring the net current from each component to the choice outputs (Fig. 2c). During trials where response 1 was chosen (mapping 1 trials), residual decision representations excited the response 1 output unit and inhibited the response 2 output unit, particularly within the choice epoch (Fig. 2d, center). During mapping 2 trials, these representations had the opposite effect. In contrast, the mean decision representations inhibited both response choices throughout the sample and delay epochs, but not the choice epoch (Fig. 2d, left). This was essential to preventing premature response choice initiation during the delay epoch (Fig. 2d, center). The contribution of the stimulus representations to response selection was negligible throughout the trial (Fig. 2d, right). Quantitatively similar results were obtained for all consecutively learned 50-problem groups in all the networks that we tested. These results demonstrate that the decision manifold constitutes the neural correlates of the task’s schema — it represents the shared temporal (mean decision) and 2-alternative (residual decision) structure of the task in an abstract form, and thereby reflects knowledge abstracted from past experiences. This predicts that the decision manifold facilitates generalization of the task structure.

### Schematic manifold embodies a representational scaffold that facilitates learning

We have shown that the schematic decision manifold is reused by, or *scaffolds* [8, 38, 39], the learned representations in subsequent problems. Moreover, this reuse is accompanied by a stark improvement in learning efficiency between the first problem and subsequent ones (Fig. 1d). To establish whether the reuse of the decision manifold plays a causal role in improving learning efficiency, we compared the learning efficiency of networks that were barred from reusing the existing decision manifold to control output responses in new problems, with networks that were allowed to do so. This method has been applied in brain-computer interface (BCI) studies to establish a causal link between monkeys’ ability to rapidly adapt to BCI readout perturbations and their reuse of existing motor cortical representations to modulate the perturbed readouts [33].

In our model, this intervention relies on the concept of a *readout subspace*. Population activity modulates an output unit’s response, only when the sum of the excitatory and inhibitory postsynaptic currents it produces at the unit are non-zero (output-potent activity, [40]). Since these currents depend on the model’s output connection weights, the output weights constrain the set of population activity levels which can modulate output unit responses. This set defines the readout subspace of population state space. Our observation that population representations within the decision subspace predominantly modulate output responses implies that the decision subspace strongly overlaps with the readout subspace. It follows that the elimination of this overlap, by appropriately altering the readout subspace, would force the development of new decision representations to modulate the output responses. This would impair the effectiveness of the representational scaffold provided by the pre-existing decision manifold in composing the learned trajectories. The observation of a concurrent learning deficit would establish a causal link between the representational scaffold and accelerated learning. For this causal intervention as well as its controls, we started by training a naive network on a single problem so that it appropriately developed overlapping readout and decision subspaces (Fig. 3a).

**Figure 3:**
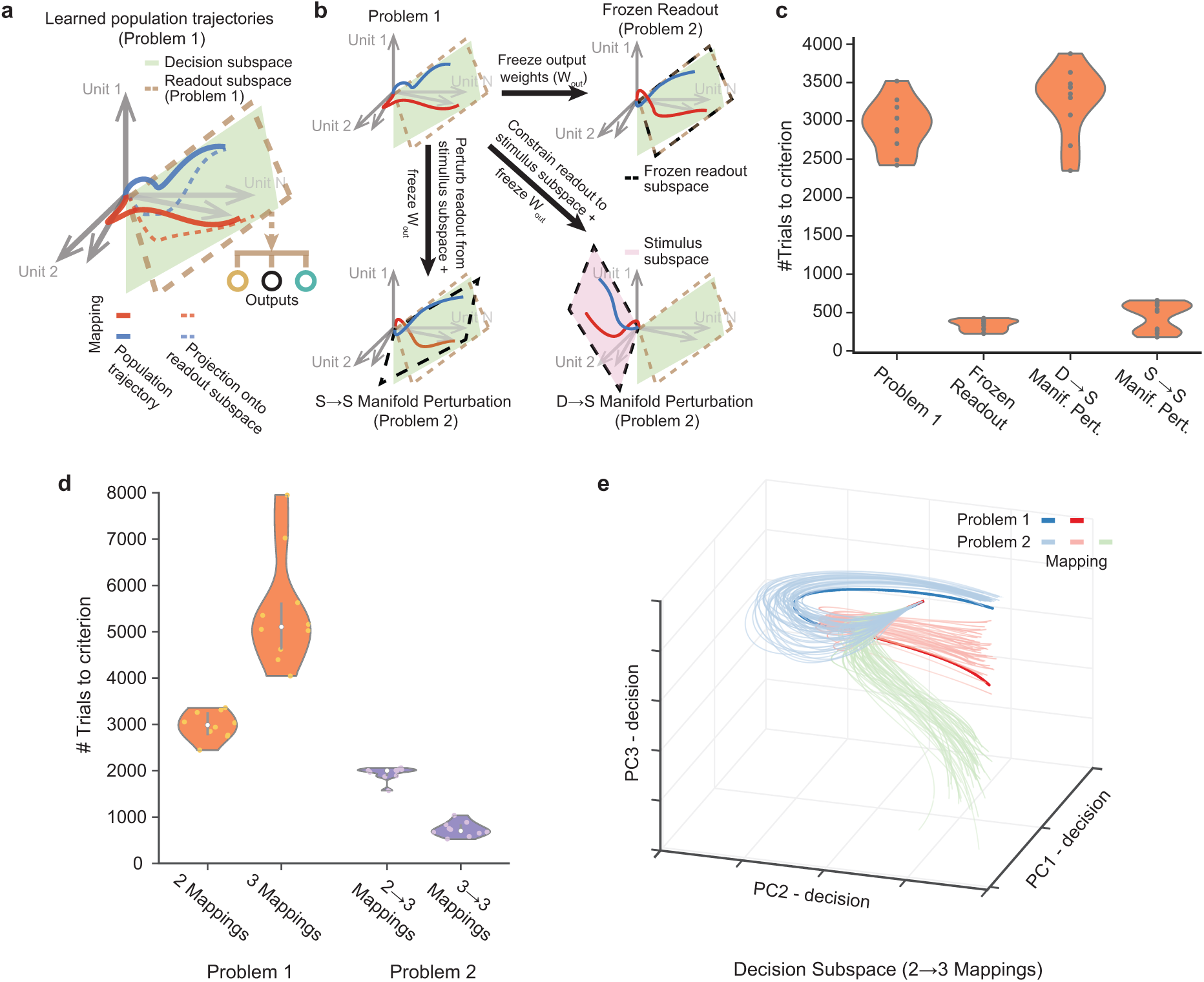
Manifold perturbations reveal that the reuse of the schematic decision manifold facilitates learning. **a.** Output responses are readout from a subspace of the population state space, which is spanned by the network’s output weights. Overlap between this readout subspace and the decision subspace enables the control of output responses by the decision representations. **b.** Illustration of the manifold perturbation interventions that assess the role of decision manifold reuse in learning. A network is trained on a single problem to establish its decision and readout subspaces (top left). It is then trained on a second problem (i) while its output weights are frozen (frozen readout, top right), (ii) after perturbing and freezing its output weights such that its readout subspace only overlaps with its stimulus subspace (D → S manifold perturbation, bottom right), or (iii) after perturbing and freezing its output weights such that the overlap between its readout and stimulus subspaces is altered (S → S manifold perturbation, bottom left). **c.** The average learning efficiency on the second problem in each of the three conditions, compared to the learning efficiency on the first problem. **d.** The learning efficiency on the first problem comprised of two versus three mappings, compared with the average learning efficiency on the second problem. The latter is a three-mapping problem and was preceded either by a two-mapping (2 → 3 mappings) or three-mapping (3 → 3 mappings) problem. **e.** Decomposed neural representations in the 2 → 3 mapping condition. Plot shows learned representations for the second problem in the first 3 principal components of the decision subspace (light), and the decision representations for the first problem projected into the same subspace (dark). Second problem decision representations are shown for 50 independently chosen stimulus sets. Trials-to-criterion on the second problem is averaged over 50 independently chosen random perturbations (**c**) / stimulus sets (**d**), and presented as the distribution of these average learning efficiencies across 10 networks with different initial conditions.

In the frozen readout condition, we then trained the network on its second problem while freezing (or preventing changes to) the output weights (Fig. 3b, top right). This helped assess whether freezing the output weights adversely affects learning efficiency. Results showed that networks exhibited a substantial improvement in learning efficiency from the first problem to the second despite frozen readouts (Fig. 3c).

In the stimulus-to-stimulus (S *→* S) manifold perturbation condition, we perturbed the output weights such that the overlap between the readout and stimulus subspaces was altered, but the overlap between the readout and decision subspaces was not (Fig. 3b, bottom left; see *Methods*). We then trained the network on its second problem with frozen output weights, which prevented the training procedure from re-aligning the readout subspace with the stimulus subspace. In these networks as well, we found a substantial improvement in the learning efficiency from the first problem to the second (Fig. 3c).

Finally, in the decision-to-stimulus (D *→* S) manifold perturbation condition, we perturbed the output weights such that the readout subspace overlapped exclusively with the stimulus subspace (Fig. 3b, bottom right). We then trained the network on its second problem with frozen output weights. This condition eliminates any overlap between the readout and decision subspaces, and compels the formation of new decision representations within the original stimulus subspace. In contrast to networks with frozen readouts and S *→* S manifold perturbations, we found that the learning efficiency of these networks was strongly impaired — they learned as slowly as naive networks learning their first problem (Fig. 3c). This demonstrates that D *→* S manifold perturbations adversely affected learning performance because the reuse of the decision manifold was impeded, and not because this was achieved by perturbing and freezing the output weights.

We further tested whether the transfer of prior knowledge could facilitate learning of problems with altered but overlapping task structure. To do so, we trained a naive network on a single problem comprised of two mappings as in Figure 3a. Then, we transitioned it to its second problem that was comprised of three mappings (i.e. three sensory stimuli mapped to three motor responses). Here too, we observed a substantial facilitation of learning performance compared to a naive network (Fig. 3d), accompanied by a reuse of the decision manifold when learning the three-mapping problem (Fig. 3e). Taken together, these results confirm that the schematic decision manifold forms a representational scaffold that facilitates the transfer of prior knowledge regarding the task’s structure to new problems, and expedites learning in the process.

### Representational reuse and synaptic plasticity differentially contribute to learning

We have shown that the reuse of existing representations to learn problems improves learning efficiency. However, learning produces fairly large changes in population activity to mediate the necessary output response corrections (Supplementary Fig. 8b). How does the emergence of such sizeable changes benefit from the reuse of existing representations? And how do the contributions of this reuse compare to those of the plasticity-induced connection weight changes? To answer these questions, we analyzed the activity changes between the beginning and end of a problem.

The neural population responds to a novel sample stimulus with a *pre-learning* trajectory in population state space (Fig. 4a right, blue curve). This trajectory evolves through time due to the repetition of the following process (equation (2)): The inputs and population activity at time *t −* 1 generate postsynaptic currents subject to the network’s recurrent and input connection weights; these currents are integrated by the network units, which advances their activity from 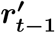 to 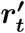 (Fig. 4a, left). In state space, this instantaneous advance in the population state is represented by a vector originating at 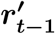 (Fig. 4a, right). Note that the direction and magnitude of this advance is state-dependent — it depends on the activity levels of the population’s units, i.e. the population state, at time *t −* 1. It also depends on the network’s connection weights. The temporal sequence of these vectors guides the evolution of the population’s activity between its initial 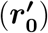 and final 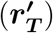 states (Fig. 4a, right, blue arrows along blue curve). Importantly, these state-dependent vectors constitute a *vector field* [41, 42] that spans the entire state space and describes the network’s dynamics (Fig. 4a, right, blue arrows tiling the space).

**Figure 4:**
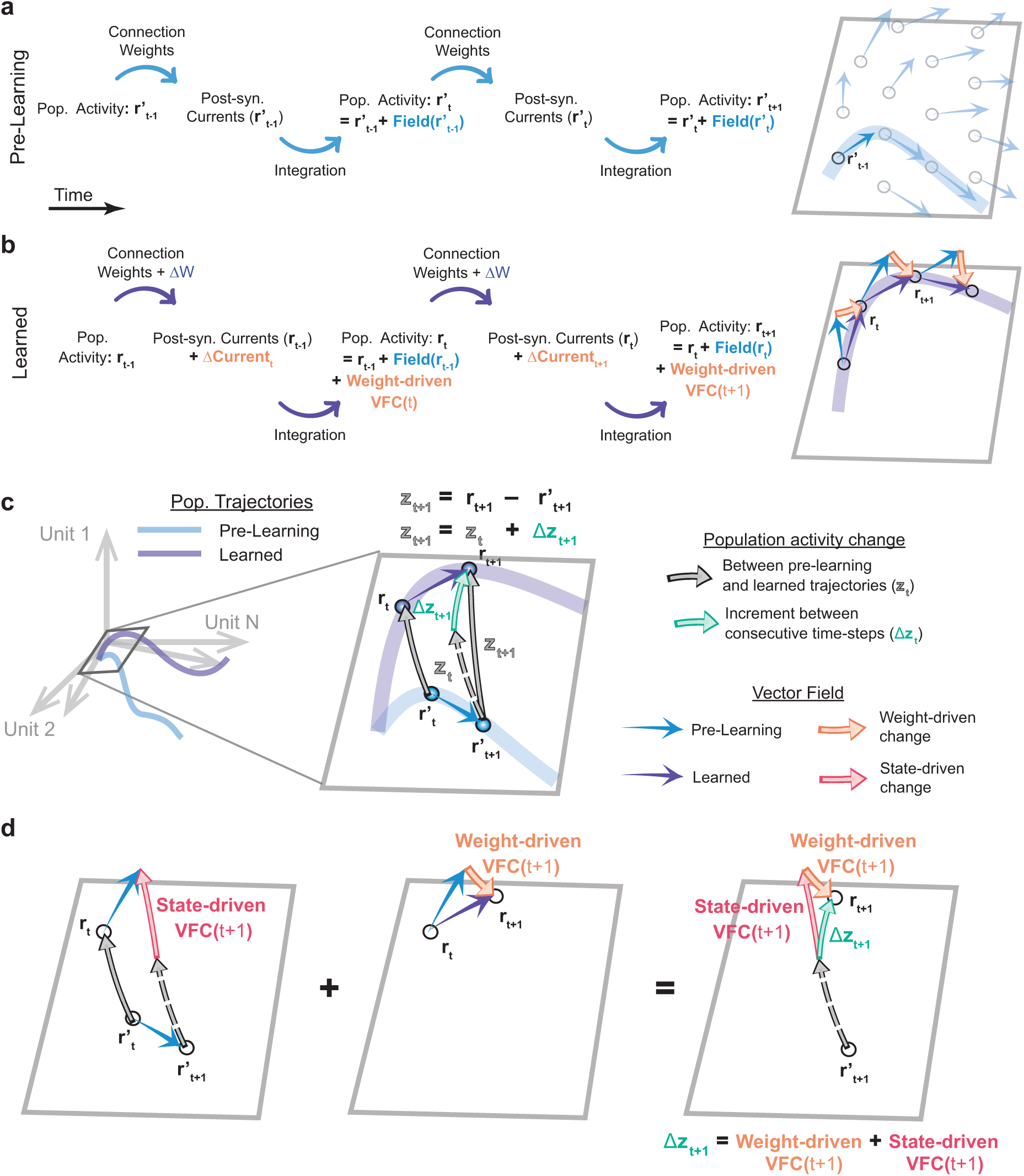
Learned trajectories emerge from vector field changes. **a-b.** Schematic describing the temporal evolution of population activity at the start (Pre-learning, **a**) and end (Learned, **b**) of a problem, with illustrations of this evolution in population state space (right). **a.** The activity advances due to the integration of net postsynaptic currents, which depend on the activity levels (or state) of the network and input units and their efferent connection weights (left). This population state-dependent advance determines a vector field that tiles state space (right, blue arrows) and guides the evolution of the population trajectory (right, blue curve). **b.** Plasticity-induced connection weight changes (ΔW) alter the postsynaptic currents (Δ**Current**), thereby altering the advance in population activity (left). The effect of this weight-driven vector field change is a continual series modifications to the vector field (right, orange arrows) that determines the evolution of the learned population trajectory (right, purple curve). **c.** The divergence of the learned trajectory from the pre-learning trajectory (**z_t+1_**, right, solid gray arrow) emerges from an accumulation of activity change increments throughout the trial (Δ**z_t+1_**, right, green arrow). **d.** Each increment is the sum of the state- and weight-driven vector field changes (left and center, pink and orange arrows, respectively). The state-driven vector field change is a result of state-dependent differences in the pre-learning vector field, specifically between learned and pre-learning population states (left, blue arrows at **r_t_** and 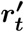, respectively). Dashed gray arrows in (c, d) represent a displaced version of the vector **z_t+1_** to help illustrate vector differences.

When the problem is learned, the population activity traverses a *learned* trajectory (Fig. 4b right, purple curve) comprised of learned population states. Since the connection weights at the end of a problem are given by the sum of the pre-learning weights and the plasticity-induced weight changes, this learned trajectory is governed by the sum of the pre-learning vector field and the changes in this field caused by the weight changes. Consequently, changes in population activity from the pre-learning to the learned trajectory are also governed by these two factors. The change in population activity from a pre-learning state 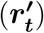 to a learned state (***r_t_***) at time *t*, ***z_t_***, is represented by a vector in state space from the former to the latter (Fig. 4c, solid gray arrows). It emerges from an accumulation of activity change increments throughout the trial (Fig. 4c, green arrow). The incremental change in population activity (Δ***z_t_*_+1_**) between times *t* and *t* + 1 derives from the pre-learning vector field (i.e. the reuse of existing representations) and the plasticity-induced changes in the vector field.

Setting aside the effect of weight changes for a moment, consider the network’s pre-learning vector field at the learned and pre-learning states. Due to its state-dependence, the pre-learning vector field may advance population activity quite differently at one state versus at the other. In this event, the activity difference between the pre-learning and learned states will change between times *t* (***z_t_***) and *t* + 1 (***z_t_*_+1_**). In state space, this change is reflected by the vector difference (Fig. 4d, left, pink arrow) between the pre-learning vector field at the two states (blue arrows), and is referred to as the *state-driven vector field change* (or state-driven VFC; referred to in *Methods* as Δ***Field_s,t_*_+1_** - equation (6)). The state-driven VFC solely depends on the pre-learning vector field (i.e. on existing representations).

The connection weight changes alter the net postsynaptic currents into the population, thereby altering how the population activity advances over time (Fig. 4b, left). In state space, this translates to a change in the vector field all along the learned trajectory (Fig. 4b, right, orange arrows), including at time *t* (Fig. 4d, center), and it is referred to as the *weight-driven vector field change* (or weight-driven VFC; referred to in *Methods* as Δ***Field_w,t_*_+1_** - equation (7)). The sum of these two types of vector field change (weight-driven and state-driven VFCs) produces the incremental change in population activity (Δ***z_t_*_+1_**) between times *t* and *t* + 1 (Fig. 4d, right; equations (4)-(5)).

Measurements revealed a substantial difference between the magnitudes of the activity changes (***z_t_***; Supplementary Fig. 8b) and the activity change increments (Δ***z_t_***; Fig. 5b). That is, large population activity changes emerge from an accumulation of relatively small change increments generated throughout the trial. We further assessed the relative contribution of the weight-driven and state-driven VFCs to the activity change increments by decomposing them (Fig. 5a; see *Methods*) into their respective components in the direction of the activity change increments (Δ***z***_∥_ - parallel component) and orthogonal to them (Δ***z_⊥_*** - orthogonal component).

**Figure 5:**
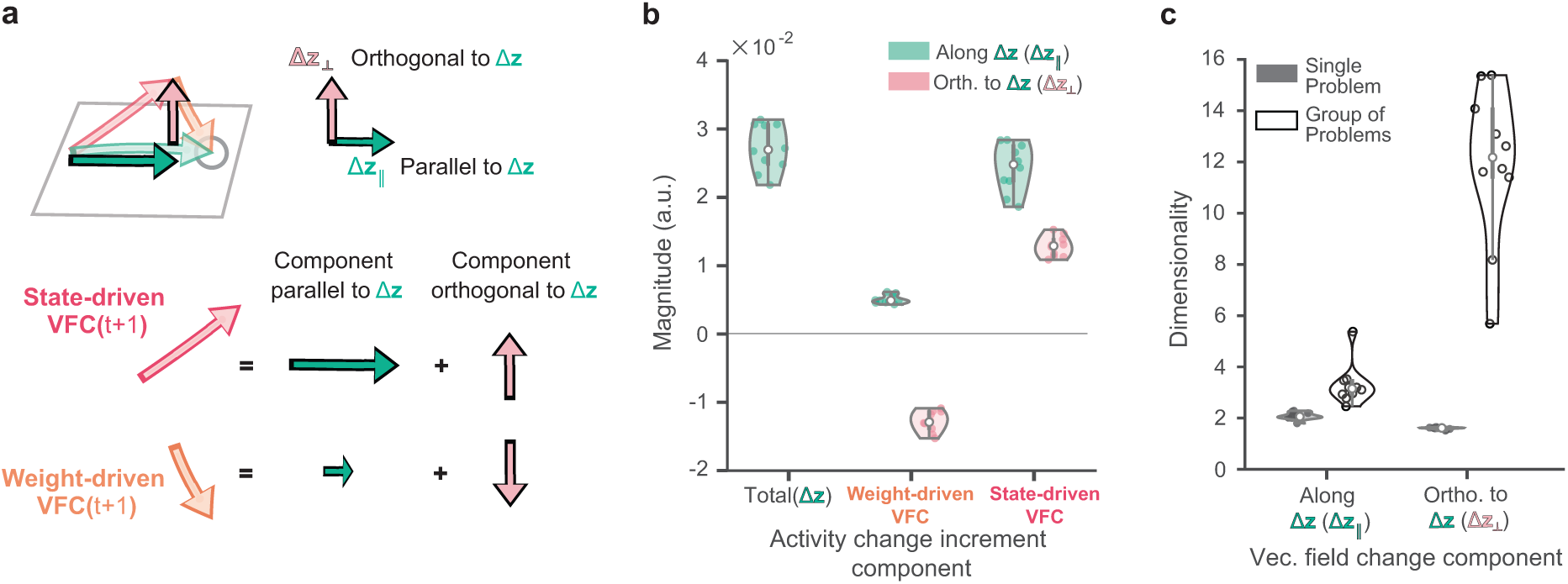
Weight- and state-driven vector field changes differentially contribute to population activity changes. **a.** The state- and weight-driven vector field changes may be decomposed into components along (or that contribute to) the activity change increment (green arrows), and components orthogonal to it (or that cancel out, pink arrows). **b.** Magnitude (L^2^- norm) of the population activity change increments, and its vector field change constituents (weight- and state-driven vector field changes) decomposed along and orthogonal to the population activity change increments. Measurements shown are the temporal mean of the magnitudes over the trial duration, averaged over both mappings of problems 2-51. **c.** Dimensionality of the vector field change components on single problems (averaged over problems 2-51) and for a group of 50 problems (problems 2-51). Plots represent distributions over 10 networks with different initial conditions.

A key observation was that the state-driven VFC’s parallel component is much larger in magnitude than the weight-driven VFC’s parallel component (Fig. 5b, green bars). Therefore, the network’s pre-learning vector field, which governs the state-driven VFC, is primarily responsible for the population activity changes. Dimensionality measurements of these parallel components revealed that they are low-dimensional not only in individual problems, but also across a group of problems (Fig. 5c). This is consistent with the structure learning hypothesis [23], which posits that efficient learning relies on changing behavior via parametric changes within a low-dimensional internal parameter space of the brain. Our results suggest that this low-dimensional internal parameter space corresponds to a low-dimensional subspace of neural population activity, which constrains how population activity may change when learning a problem.

Measurements also showed that the weight-driven VFC’s orthogonal component is much larger in magnitude than its parallel component, and that it is equal in magnitude but opposite in direction to its state-driven counterpart and nullifies it as a result (Fig. 5b, pink bars). These orthogonal components of the weight- and state-driven VFCs are also low-dimensional on individual problems, but are high-dimensional across a group of problems (Fig. 5c). Moreover, they largely span directions along which the existing representations do not typically covary (Supplementary Fig. 9a). These results imply that novel sample stimuli interact with the existing representations when mapped onto them, in a manner that elicits uncharacteristic population responses. They also reveal that the existing representations can be sensitive (i.e. not entirely invariant) to the sample stimuli that are mapped onto them. The weight-driven VFC emerges primarily to impede such interactions and thereby prevent changes to the existing representations.

To summarize, our analysis of the population activity changes between the start and end of problem learning revealed that: (i) large changes emerge over the trial timecourse from the accumulation of a sequence of small local changes along the learned trajectory; (ii) these changes are low-dimensional and stem primarily from reusing the network’s pre-learning vector field to shape the learned trajectory, thus elucidating the relative contribution of representational reuse to learning; (iii) the pre-existing representations are not entirely invariant to having novel sample stimuli mapped onto them, and can undergo uncharacteristic modifications in the process. Connection weight changes emerge largely to prevent such modifications.

### Magnitude of recurrent weight changes determines learning efficiency

Next, we examined why learning efficiency is enhanced by representational reuse, by exploring how learning efficiency is impacted by the connection weight changes. Before we could do so, it was important to evaluate the relative contribution of input versus recurrent weight changes to learning. In supplementary note 1.1, we show that the model learns via recurrent weight changes — these changes predominantly determine the weight-driven VFC — as it is more efficient to do so. Moreover, measurements showed that the magnitude of recurrent weight changes in a problem largely explains the number of trials expended in learning it (Fig. 6a, left; coefficient of determination *R*^2^=0.7). This relationship was robustly observed across all 10 networks that were tested (Fig. 6a, right) and is consistent with analytical bounds relating the magnitude of connection weight changes and sample efficiency in deep neural networks [43, 44].

**Figure 6:**
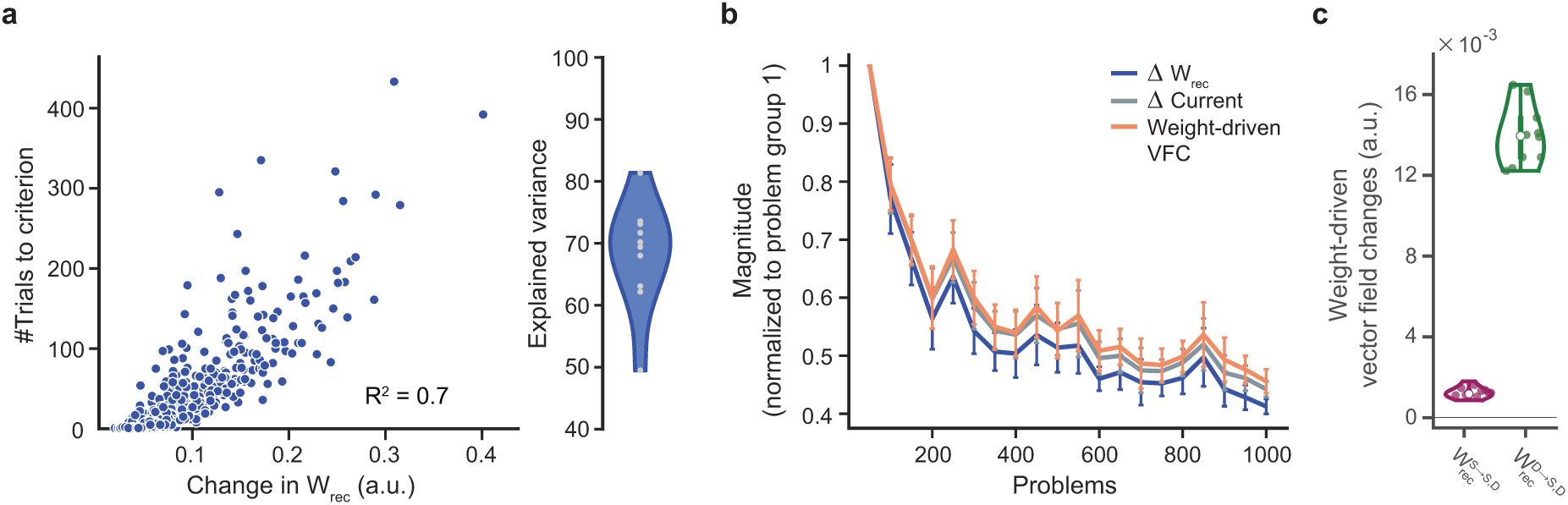
The magnitude of recurrent weight changes explains both the magnitude of the weight-driven VFC and the number of trials to learn a problem. **a.** The magnitude of the plasticity-induced recurrent connection weight changes explains a majority of the variance in the number of trials to learn problems (left). This relationship was robustly observed across ten networks with different initial conditions (right). **b.** The magnitude of recurrent weight (blue), postsynaptic current (gray) and weight-driven vector field (orange) changes, averaged in groups of 50 non-overlapping and consecutively learned problems. Each quantity has been normalized by its corresponding value for the first problem group. All quantities decrease exponentially with the number of previously learned problems. **c.** Approximate contribution of presynaptic population activity in the stimulus versus decision subspace to the weight-driven vector field change, averaged over problems 2-51. The magnitudes (L^2^-norm) of the change in the postsynaptic currents and vector field represent their temporal mean over the entire trial duration, averaged over both mappings in each problem. The magnitude of recurrent weight changes was measured by their Frobenius norm. Plot (b) (plot (c)) reflects mean values (the distribution) over 10 networks with different initial conditions, and the error bars indicate their standard errors.

In light of this observation and the exponential decrease in the trials-to-criterion across problems, we hypothesized that the magnitude of recurrent weight changes should also decrease exponentially over the sequence of learned problems. We further posited that the recurrent weight change magnitudes should be proportional to the postsynaptic current change and the weight-driven VFC magnitudes, since these three quantities are directly related to each other. Consequently, we expected that the magnitudes of the postsynaptic current changes and the weight-driven VFC would also decrease exponentially. Figure 6b confirms that the magnitude of these three quantities decreases exponentially as a function of the number of previously learned problems. Therefore, the progressive improvement in the model’s learning efficiency is explained by a similar decrease in the magnitudes of the recurrent weight changes and weight-driven VFC required to learn problems.

We can now explain why the reuse of existing representations markedly improves learning efficiency (Fig. 3). Networks with D *→* S manifold perturbations are compelled to develop new representations of the task’s structure beyond the original decision sub-space, and aligned with their perturbed readout subspace (Fig. 3b, bottom right). In other words, the structure and location, in state space, of the target trajectories are largely constrained by the arbitrarily altered output weights. However, the vector field along such an arbitrarily constrained target trajectory is most likely misaligned relative to the vector field required to support it (Supplementary Fig. 7a, right, purple versus blue arrows along learned trajectory). Consequently, it is unlikely to roughly advance population activity along the target trajectory, as it does in unperturbed networks (Supplementary Fig. 7a, left). Measurements comparing the magnitude of the weight-driven VFC in unperturbed and perturbed networks confirms that the vector field in perturbed networks undergo drastic re-organization in comparison to unperturbed networks (Supplementary Fig. 7b, right), so that they may shape the trajectories that will re-encode the task’s structure (large orange arrows, Supplementary Fig. 7a). This explains the impairment in learning efficiency following D *→* S manifold perturbations and demonstrates the merits of learning via representational reuse — it is this reuse of existing representations that limits the requisite weight changes (Supplementary Fig. 7b, left), and thereby improves the learning efficiency.

In supplementary note 1.2, we explore the interactions between stimulus and decision representations during trial performance and learning. We found that the stimulus and decision representations exhibit reciprocal interactions to sustain each other through the trial, and that the weight-driven VFC largely prevents uncharacteristic changes in the existing stimulus representations (rather than existing decision representations). In addition, we assessed whether the weight-driven VFC is modulated more strongly by pre-synaptic population activity in the stimulus or decision subspace. A comparison of their approximate contributions to the weight-driven VFC revealed that it relies almost entirely on decision representations (Fig. 6c, Supplementary Fig. 8d). This is likely due the the fact that the decision representations are larger in magnitude than the stimulus representations. These results reveal a second form of representational scaffolding, wherein the decision representations scaffold the formation of the weight-driven VFC.

### Accumulation of weight changes across problems progressively improves learning efficiency

In agreement with Harlow’s learning-to-learn experiments, our model exhibits a progressive improvement in learning efficiency spanning a few hundred problems (Fig. 1). This improvement is explained by a progressive decrease in the magnitude of the weight changes and weight-driven VFC per problem (Fig. 6a-b). Since the weight-driven VFC in a problem primarily prevents distortions to existing representations during learning (Fig. 5b), a progressive decrease in its magnitude amounts to a progressive improvement in the invariance of the existing representations to having novel stimuli mapped onto them. However, the source of this improvement is yet undetermined: what causes it in the absence of an explicit meta-learning mechanism, and how does the network’s accumulation of learning experience over problems relate to its emergence? We hypothesized that the accumulation of weight changes over earlier problems facilitates learning in future problems. That is, weight changes elicited while learning problems *p − k* (for 1 *≤ k ≤ p −* 2) cumulatively alter the vector field such that they suppress the weight-driven VFC required to learn problem *p* (Supplementary Fig. 10a, top; see *Methods*). More generally, as problems are learned, their respective weight-driven VFCs accumulate to produce a *cumulative* vector field change (or cumulative VFC) which suppresses the weight-driven VFC required to learn subsequent problems. This progressively improves representational invariance and thereby accelerates learning.

To test this hypothesis, for each problem *p*, we measured the magnitudes of its weight-driven VFC plus the cumulative VFC along its learned trajectory due to the accumulation of weight changes over the sequence of problems that precede it, i.e. from problem *p* − 1 (relative problem −1) to problem 2 (relative problem 2 *−p*). Figure 7a summarizes these measurements across many problems *p* grouped by their learning-to-learn stage, i.e. the number of problems they are preceded by. Here, we focused on the magnitude along each problem’s orthogonal weight-driven VFC component (Δ***z_⊥_***), because it dominates the total weight-driven VFC in problems at each learning-to-learn stage (Supplementary Fig. 9b). The results indeed show that at each stage, learning earlier problems cumulatively suppresses the weight-driven VFC required in subsequent problems. We further found that this is predominantly due to an accumulation of recurrent weight changes (Supplementary Fig. 9c). These findings confirmed our hypothesis — the accumulation of weight changes over problems progressively improves representational invariance and therefore learning efficiency. Moreover, they imply that the cumulative change along the orthogonal weight-driven VFC component of problems imposes a learning efficiency bottleneck.

**Figure 7:**
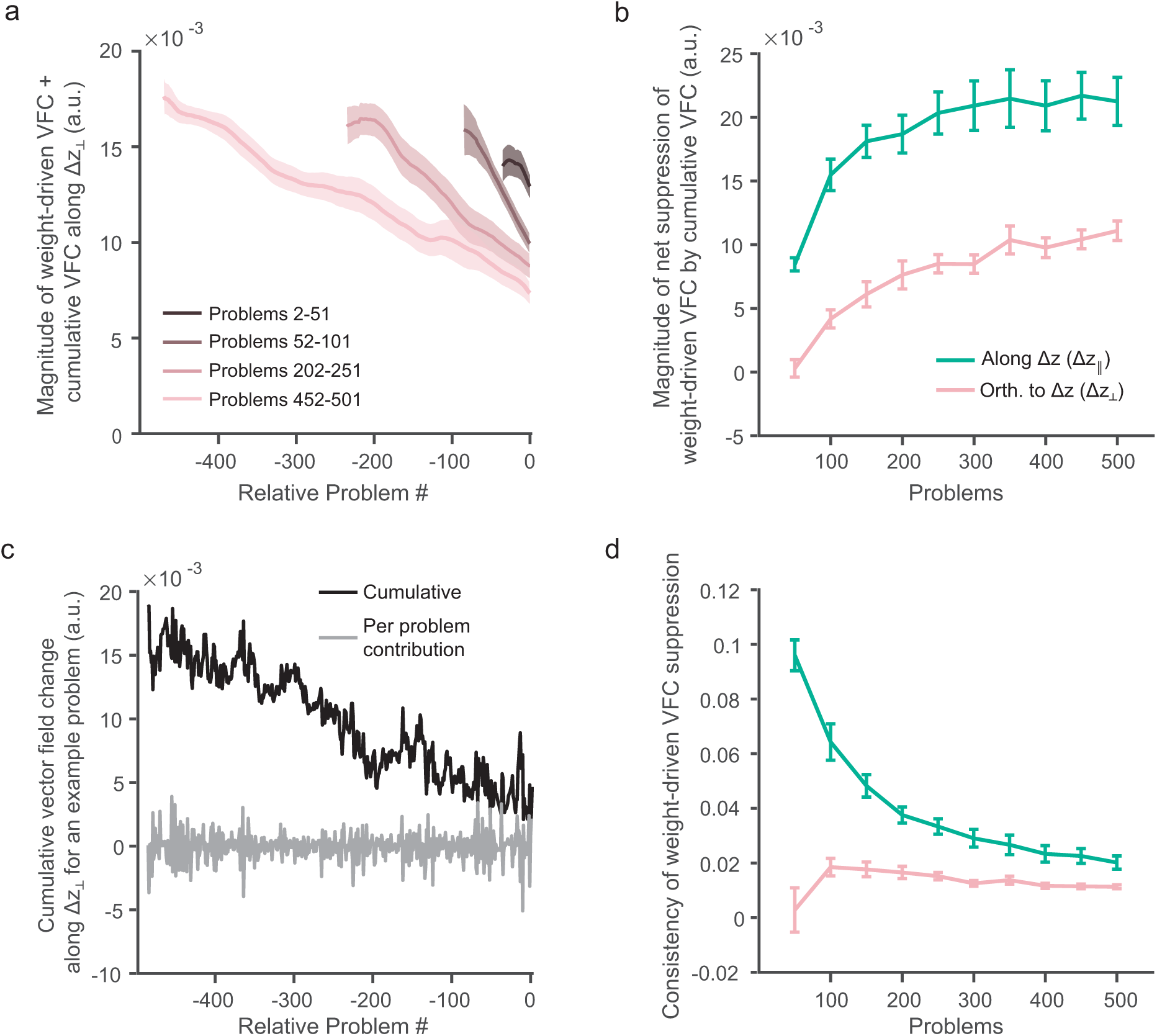
Accumulation of weight changes progressively improves invariance of existing representations to learning. **a.** Magnitude of vector field change along the learned trajectory for a problem p due to the accumulation of (i) the weight changes in problem p (W^p^− W^p−1^, relative problem = 0; weight-driven vector field change), and (ii) weight changes in each of the earlier problems, proceeding backwards to problem 2 (W^p^ − W^p−k−1^ for 1 ≤ k ≤ p − 2, relative problem −k; cumulative vector field change contributions). The curve for each problem measures the magnitude of change in the direction of its orthogonal weight-driven VFC component, smoothed with a 30-problem moving average filter. Plot summarizes the measurements for problems in 4 problem groups at different stages of learning-to-learn, and demonstrates the suppressive effect of the cumulative vector field change at each stage. **b.** Magnitude of net suppression for each problem p, due to the net weight changes between the start of problems 2 and p (W^p−1^ − W^1^), summarized in 50-problem groups. **c.** Magnitudes of the cumulative VFC and cumulative VFC contributions by individual problems along an example problem’s orthogonal weight-driven VFC component. **d.** Ratio of the net suppression magnitude to the sum of magnitudes of the cumulative VFC contributions, summarized as in (b). This measures how consistently suppressive the cumulative VFC contributions for a problem are. Measures in (b, d) are presented separately for vector field changes along the parallel (green) and orthogonal (pink) weight-driven VFC components. Magnitudes shown are the temporal mean of the unsigned (L^1^-norm; a) and signed (b, c) projections onto the parallel / orthogonal weight-driven VFC components, averaged over both mappings in a problem. Plots in (a, b and d) reflect mean values over 10 networks with different initial conditions, and shading/error bars indicate standard errors.

Surprisingly, even though the network expends many more trials on learning early problems, the approximate linearity of the curves in figure 7a indicates that early- and late-learned problems produce similar-sized contributions to the cumulative VFCs. Indeed, measurements showed that the per-trial cumulative VFC contributions by late-learned problems are larger than those by early-learned problems (Supplementary Fig. 11a). This demonstrates that with experience, the model learns to contribute to the learning efficiency of future problems in an increasingly efficient manner.

Figure 7a further demonstrates that the weight-driven VFC in a problem depends on its net suppression by the preceding problems, i.e. the sum of the suppressive cumulative VFC contributions (and enhancing cumulative VFC contributions when they increase the requisite weight-driven VFC) by the weight changes in each preceding problem going back to problem 2 (Supplementary Fig. 10b, left; see *Methods*). A larger net suppression produces a smaller weight-driven VFC. Since the weight-driven VFC decays exponentially with the number of preceding problems (Fig. 6b), we posited that the net suppression must similarly increase with it. Measurements of the net suppression along the orthogonal and parallel weight-driven VFC components of problems confirmed this (Fig. 7b). The net suppression mirrors the exponential decay in the weight-driven VFC (see *Methods*) — it rapidly increases across problems at the early stages of learning-to-learn, which produces a rapid decrease in their weight-driven VFCs; it gradually plateaus for later problems, which explains the plateauing of their weight-driven VFCs. The results also showed that the net suppression is weaker along the orthogonal components than along the parallel components, which explains why the learning efficiency bottleneck develops along the orthogonal components.

Figure 7b also revealed that the net suppression of a problem’s weight-driven VFC is not linearly related to the number of problems that precede it. This indicates that the rate at which the cumulative VFC contributions suppress a problem’s weight-driven VFC depends on its learning-to-learn stage. We reasoned that slow (quick) suppression must be due to smaller (larger) contributions by the weight changes in preceding problems. Therefore, we expected that the sum of the magnitudes of these contributions would be small (large) for problems whose weight-driven VFC is suppressed slowly (quickly), and the progression of this sum over the learning-to-learn stages would resemble the net suppression magnitudes (Fig. 7b). Instead, we found that the sum increases in a largely linear fashion for the cumulative VFC contributions along both the parallel and orthogonal weight-driven VFC components (Supplementary Fig. 11b). This indicated that *(i)* the weight-driven VFC of problems at different learning-to-learn stages are altered to a similar extent by weight changes in the problems that precede them, and *(ii)* the sum of the magnitudes of these cumulative VFC contributions reflects the number of problems that they accumulate over, but not the differences in their suppression rates. Taken together with the results in figure 7b, this reveals a surprising result: although the cumulative VFC contributions are similar-sized, their suppressive effect on a future problem’s weight-driven VFC depends on its learning-to-learn stage.

Interestingly, along both the parallel and orthogonal weight-driven VFC components, we observed that the sum of contribution magnitudes (Supplementary Fig. 11b) is orders of magnitude larger than the net suppression magnitudes (Fig. 7b). In other words, relatively large cumulative VFC contributions from earlier problems cumulatively suppress an ensuing problem’s weight-driven VFC by relatively small amounts. Based on this observation, we concluded that the effect of weight changes in individual problems on a given future problem’s weight-driven VFC are largely inconsistent with each other; some cumulative VFC contributions suppress the problem’s weight-driven VFC while others enhance it (Supplementary Fig. 10b, right). In fact, the ratio of the net suppression magnitude to the sum of contribution magnitudes quantifies this consistency; a value of 1 would indicate that the problem’s weight-driven VFC is suppressed by the weight changes in each of its preceding problems, and a value of 0 would indicate that the cumulative VFC contributions are maximally inconsistent - the enhancing and suppressing contributions nullify each other resulting in no net change. For problems at all learning-to-learn stages, we found that this ratio was closer to 0 (Fig. 7d).

These results depict the suppressive effect of the accumulating weight changes on each future problem’s weight-driven VFC as a stochastic process — cumulative VFC contributions by individual problems stochastically enhance or suppress the future problem’s weight-driven VFC. However, they collectively exhibit a weak bias towards consistently suppressing it (values in Fig. 7d are above zero). The cumulative effect of this weakly suppressive bias is a small yet significant suppression of the weight-driven VFC. We illustrate this process along the orthogonal weight-driven VFC component of an example problem. Since the orthogonal weight-driven VFC components in a problem are roughly one-dimensional (Fig. 5c), the stochastic process is one-dimensional as well. The individual cumulative VFC contributions by earlier problems towards suppressing the weight-driven VFC along this dimension in the example problem (grey curve) fluctuate between positive (suppressing) and negative (enhancing) values (Fig. 7c). However, due to their weakly suppressive bias, these contributions cumulatively produce a large net suppression of the problem’s weight-driven VFC (black curve).

Figure 7d further shows that the cumulative VFC contributions are more inconsistent along the orthogonal weight-driven VFC components than along the parallel components, indicating that their suppressive bias is weaker along the orthogonal components. This explains the weaker net suppression along the orthogonal components (Fig. 7b), and why it imposes a learning efficiency bottleneck. Crucially, figure 7d demonstrates that the weight-driven VFC of problems at different learning-to-learn stages are suppressed at different rates due to differences in the consistency with which the weight changes in the preceding problems suppress them (Supplementary Fig. 10b). That is, the exponential decay in the weight-driven VFC magnitudes stems from a modulation of the suppressive bias in the cumulative VFC contributions. The exponential decay in the weight-driven VFC magnitudes is largely caused by an exponential decay in magnitude of their orthogonal components (Supplementary Fig. 9b). The bias in the cumulative VFC contributions to suppress the orthogonal weight-driven VFC component in early stage problems rapidly increases (Fig. 7d). This rapidly increases their net suppression (Fig. 7b), which rapidly decreases the weight-driven VFC required to learn them. Subsequent problems follow a more prolonged accumulation of weight changes (because they are preceded by more problems), albeit with a weakened bias to suppress their orthogonal weight-driven VFC components. This results in the plateauing of their net suppression, and therefore of their weight-driven VFCs.

To summarize, our results identify a novel neural mechanism of accumulating learning experience to progressively improve learning efficiency, despite the absence of a meta-learning mechanism. It relies on the accumulation of connection weight changes over learned problems to suppress the weight-driven VFC required to learn subsequent problems and thus accelerate their learning. The model progressively accelerates learning via, (*i*) a gradual improvement in the efficiency with which weight changes contribute to the suppression of the weight-driven VFC in future problems (Fig. 7b), and (*ii*) a modulation of how consistently suppressive these contributions are (Fig. 7e). Moreover, the fact that the weight-driven VFC primarily prevents uncharacteristic representational changes from developing when novel sample stimuli are mapped onto existing representations (Fig. 5) helps elucidate the objective of this learning-to-learn mechanism: the accumulation of weight changes over early problems improves the invariance of the existing representations to having novel sample stimuli mapped onto them. This refines the model’s ability to learn via representational reuse and elicits learning-to-learn.

## Discussion

New information is far easier to learn when it is contextualized by prior knowledge. This process is thought to be facilitated by the instantiation of schemas [3, 4], which are hypothesized to correspond to neocortically encoded knowledge structures. Learning-to-learn is a constructive consequence of the reciprocal influence between learning and schema tuning, whereby schema instantiation facilitates learning, and the assimilation of learned information into the schema improves its ability to facilitate future learning. To elucidate the underlying neurobiological basis, in this work we trained an RNN model on a series of sensorimotor mapping problems, without any meta-learning. Our main findings are threefold. First, the network model exhibits accelerated learning that is quantified by an exponential time course, with a characteristic time constant and a plateau. Interestingly, this model prediction appears to be supported by an ongoing experiment where monkeys displayed an exponential learning-to-learn time course while solving a series of arbitrary sensorimotor mapping problems [36]. Second, schema formation corresponds to the formation of a low-dimensional subspace of neural population activity, thereby bridging a psychological concept with a neural circuit mechanism. Third, rather than weight changes *per se*, it is imperative to examine weight driven changes of the vector field in order to understand the behavior of a recurrent neural network as a dynamical system. These new insights can be used to guide the analysis of neurophysiological data from behaving animals during learning-to-learn.

Our work revealed that learning-to-learn is a process with three timescales (Fig. 8). The fastest timescale governs the evolution of population activity over a single trial. Sub-space decomposition of this activity showed that it encodes three latent variables. First, a mean decision component which is analogous to the condition-independent component identified in prefrontal and motor cortical activity [37, 45] — it encodes temporal aspects of the task in a trial-condition invariant manner, and explains most of the variance in population activity. Second, a residual decision component that encodes decisions and response choices. And third, a problem stimulus representation. The first two components collectively constitute low-dimensional decision representations that control fixation and response choices.

**Figure 8:**
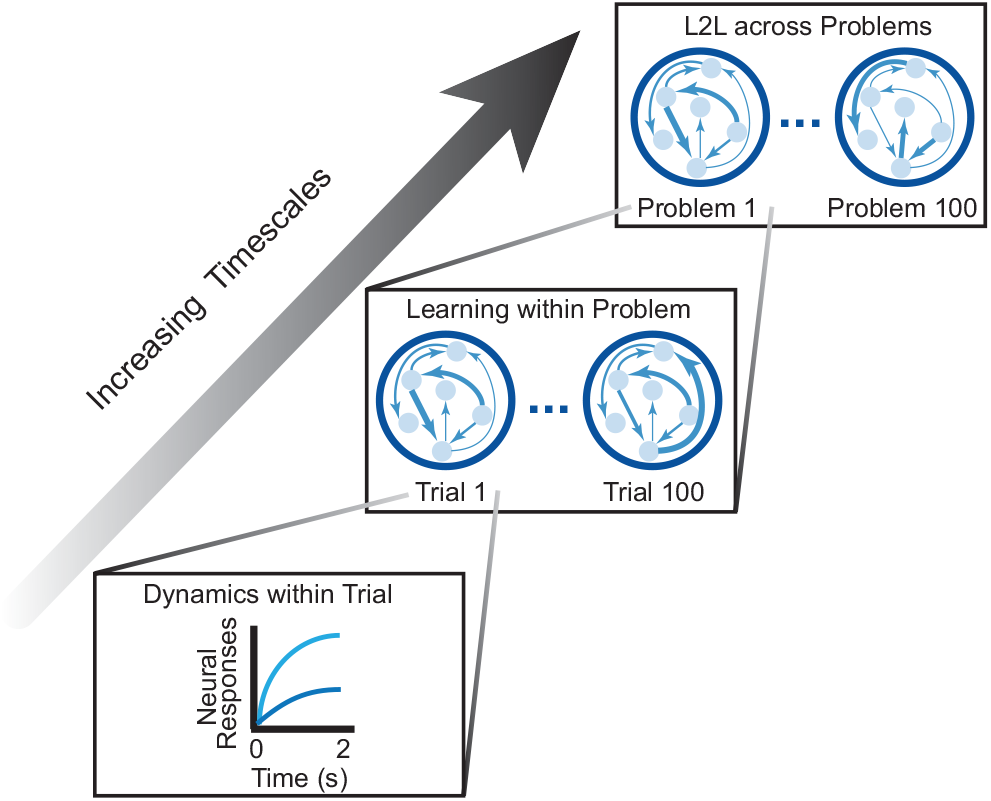
Learning-to-learn is a process with three timescales. The fastest timescale (bottom) governs the neural dynamics within a trial which drive output responses. The intermediate timescale (middle) governs the learning dynamics across trials within a problem; it ultimately produces the requisite weight-driven vector field change which results in problem being learned. The slowest timescale (top) governs the dynamics of learning-to-learn across problems; it ultimately improves the invariance of existing representations to learning new problems which results in asymptotic learning efficiency.

We found that these decision representations are shared across problems in an abstract form: the model reuses them to contextualize its neural and output responses to new sample stimuli, and to generalize from previous solutions to newer ones. Analysis of the model’s learning with a manifold perturbation intervention showed that this reuse of the decision representations causes a stark improvement in learning efficiency. These results demonstrate that the network not only abstracts commonalities across problems, but also exploits them to facilitate learning [4, 23, 46]. Therefore, the abstract decision representations constitute the neural basis of a sensorimotor mapping schema [4, 17]. It is noteworthy that the abstraction of task variable- and task structure-encoding neural representations and their reuse in consecutively learned association problems has indeed been observed in the prefrontal cortex and hippocampus [10, 21, 22].

The intermediate timescale governs the process of learning, and spans the trials between the beginning and end of learning a single problem (Fig. 8). We studied learning with a novel measure of how connection weight changes (which model the effects of long-term synaptic plasticity or LTP) influence population activity in an RNN — the weight-driven vector field change. Our results demonstrated that this measure is more informative and accurate at assessing the effects of the connection weight changes, than direct measurements of the weight changes: *(i)* it dissociates the contributions of the changes in different sets of connection weights more accurately than directly comparing their magnitudes; *(ii)* its assessments are more interpretable as they directly relate to the population activity; and *(iii)* it isolates the contributions of the initial weights and the weight changes to the learning-induced changes in population activity. For these reasons, these techniques contribute to a growing set of methods that aim to overcome the challenges of interpretability and explainability in RNNs [47, 48], which hinder their adoption in neuroscience. In our analysis, these techniques were instrumental in identifying *(i)* why reusing existing representations improves learning efficiency, *(ii)* the relative contributions of this reuse versus the connection weight changes to learning, and *(iii)* the mechanism underlying learning-to-learn.

Our analysis of the change in population activity that emerges over the timespan of learning identified two forms of schematic scaffolding. First, the reuse of existing schematic representations is primarily responsible for these activity changes. The reuse avoids the formation of new task-encoding representations, which substantially reduces the weight changes required to learn a problem. This dramatically accelerates its learning. Second, the weight-driven VFC is largely modulated by the schematic representations. Moreover, the primary effect of the weight-driven VFC is to prevent the unwarranted changes to existing representations which develop when novel sample stimuli are mapped onto and interact with them.

In the training RNN framework, the network is initialized with random weights, as a blank slate. In contrast, developmental experience shapes how new information is encoded even in the brain of a task-naive animal. This confounds direct comparisons between the use of a learning algorithm and a known biological plasticity rule. Nevertheless, our findings regarding the benefits of representational reuse do not directly depend on the learning algorithm we used, and may well be conserved under biologically plausible learning rules. Moreover, since our analysis techniques are independent of the underlying learning rules, they offer an approach to study learning and the properties of schema formation and reuse in models with biologically plausible learning rules. Our model further assumes that following schema formation, new problems continue to be learned via LTP. Indeed, rapid learning of novel schema-consistent paired associates was found to be prefrontal NMDA-receptor dependent in rodents [14], suggesting that Hebbian neocortical synaptic plasticity is likely involved in schema-facilitated learning. However, the role of other forms of plasticity, such as intrinsic [49] and behavioral timescale [50] plasticity, has not been experimentally precluded. Further computational and experimental studies are required to determine their relative roles in this process.

At the slowest timescale, several problems are learned in succession with progressively improving efficiency, until asymptotic learning efficiency is realized (Fig. 8). This is the timescale of learning-to-learn. We showed that, consistent with macaque monkeys’ behavior [29, 36], our model’s trials-to-criterion performance is well-characterized by a decaying exponential function, which asymptotes at roughly 20 trials per problem. Consequently, our model suggests that learning-to-learn can emerge in animal models in the absence of explicit meta-learning.

However, the brain may adopt one or some of the many meta-learning approaches proposed in computational neuroscience and deep learning to facilitate learning across problems. In general, meta-learning may be conceptualized as a bi-level optimization, wherein the inner-loop spans trials of a single problem over which the model’s parameters are updated to improve performance accuracy, and the outer-loop spans problems with shared task structure over which learning parameters are updated to optimize learning efficiently [51]. Biologically plausible outer-loop mechanisms include meta-plasticity of neuro-modulatory inputs to refine synaptic plasticity rates [52] and meta-plasticity of regulatory states governing synaptic plasticity to improve learning efficiency [27, 53]. A recently proposed model-based learning algorithm [25] may also be understood within this framework, wherein the outer-loop is composed of a model-free reinforcement learning algorithm to learn a task model and a corresponding model-based learning policy for problems with shared task structure, and the inner-loop is comprised of an implementation of this model-based policy via a neural population-level integration of choices and their outcomes on earlier trials to improve the accuracy of choices on the current and upcoming trials. The task model and learning policy are learned at the outer-loop by maximizing the sum of rewards over a sequence of trials of the problem learned in the inner-loop. While our approach has no explicit outer-loop mechanism, it most closely resembles the Reptile meta-learning algorithm [54]. Reptile optimizes a network model’s initial connection weights towards achieving few-shot learning of problems with shared task structure. However, our approach is different from Reptile in important ways: *(i)* Reptile’s inner-loop learns a problem for a fixed, small number of trials, rather than until the problem is fully learned, and its few-shot learning ability is quite sensitive to this number. *(ii)* Reptile’s estimate of the optimal initial connection weights is an exponential moving average of the weights learned at the end of each inner-loop, rather than the weights learned at the end of the most recent inner-loop. Its few-shot learning ability is also sensitive to the weighting factor in this exponential moving average. *(iii)* Consequently, Reptile must maintain two sets of connection weights, its current estimate of the optimal initial weights and its estimate of the update from the inner-loop iteration. Moreover, it does not specify a biological mechanism to maintain the pair of weight estimates. *(iv)* After meta-learning, Reptile learns each new problem starting from the fixed, meta-learned set of initial connection weights. Instead, our approach continues to accumulate weight changes across problems indefinitely. While several meta-learning mechanisms have been proposed by computational and deep learning studies, further study is required to identify their neural substrates and evaluate their role in learning-to-learn.

We have identified a novel mechanism for learning-to-learn, which relies on the accumulation of weight changes over learned problems to progressively improve the invariance of the existing representations to being reused with novel sample stimuli. An increase in this invariance suppresses the weight-driven VFCs required to learn new problems which accelerates their learning. Interestingly, we found that these cumulative improvements are stochastic in nature — the exponential improvement in learning efficiency stems from a modulation of the bias in this stochastic process to suppress the weight-driven VFCs in future problems.

We also found evidence in support of the structure learning hypothesis, which posits that improvements in learning efficiency are achieved by restricting the extent of parametric behavioral exploration while learning a problem [23]. It has been suggested that exploration in the space of network connection weights directly controls behavioral exploration, such that learning efficiency improves due to a progressive narrowing of the effective control space to the most task-relevant low-dimensional subspace of the space of connection weights. Instead, we found that behavioral changes are directly controlled by population activity changes within a low-dimensional subspace of population state space. In addition, connection weight changes emerge primarily to restrain activity changes to this low-dimensional subspace. Learning-to-learn derives from a progressive improvement in the model’s inherent ability to do so (i.e. without large connection weight changes) when novel stimuli are mapped onto existing representations, rather than a progressive decrease in the dimensionality of the space within which population activity or connection weights change.

Note that our results differentiate between schema-facilitated learning and structure learning. While a schema and an associated behavioral control space can emerge within a low-dimensional subspace of population state space even after the first problem is learned, structure learning proceeds thereafter and involves learning to use this control space efficiently (i.e. without large connection weight changes). Recent work has demonstrated the re-use of schematic prefrontal representations in rodents learning a series of odor-response sequence problems [10]. However, the authors did not observe an acceleration in learning. We propose that this may be explained by the presence of schema-facilitated learning, but an absence of structure learning.

Crucially, our results offer experimentally verifiable predictions. First, the sensorimotor mapping schema is encoded by low-dimensional neural representations which are shared across problems, and explain a majority of the variance in population activity. They encode shared task variables including the task’s temporal structure and the available choices. Second, the reuse of these representations to learn new problems causes a speedup in learning; preventing this reuse with recently developed BMI interventions [33] should produce pronounced learning deficits. Third, population activity may undergo large changes between the beginning and end of problem learning. However, across problems, these changes are restricted to a low-dimensional subspace of the activity. Fourth, the number of trials to learn a problem decreases exponentially as a function of the number of previously learned problems. Taken together, our results shed insights into the neural substrate of a sensorimotor mapping schema, the reason for which its reuse markedly improves learning efficiency, and the neural mechanisms of structure learning that gives rise to learning-to-learn. In doing so, they elucidate the neural mechanisms of learning-to-learn and present novel techniques to analyze learning-to-learn in RNNs.

## Acknowledgements

We thank A.L. Fairhall, I. Skelin, J.J. Lin, B. Doiron, G.R. Yang, N.Y. Masse, U.P. Obilinovic, L.Y. Tian, D.V. Buonomano and J. Jaramillo for fruitful discussions; and Y. Liu, K. Berlemont, A. Battista and P. Theodoni for critical comments on the manuscript. This work was supported by the National Institute of Health U-19 program grant no. 5U19NS107609-03 and the Office of Naval Research grant no. N00014- 17-1-2041.

## Data and Code Availability

All training and analysis codes will be available at publication on GitHub (https://github.com/xjwanglab). We will also provide data files in Python and Matlab readable formats for further analyses. Pre-trained networks will be stored in a Google Drive folder with its link provided on the same GitHub repository.

## Methods

### Recurrent Neural Network Model (Fig. 1)

The RNN model comprises a fully-connected population of *N* firing rate units with firing rates ***r***, receiving inputs from *N_in_* input units with firing rates ***u***. Firing rates of the network units follow the dynamical equation

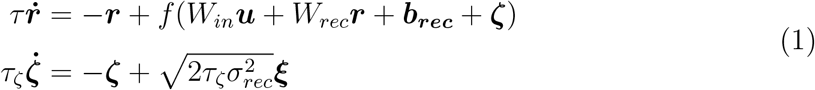

which expresses the leaky and non-linear integration of input (*W_in_****u***) and recurrent (*W_rec_**r***) currents. *W_in_* (*W_rec_*) is an *N×N_in_* (*N×N*) matrix of input (recurrent) connection weights, and *τ* = 100 ms is the integration time-constant that characterizes the slow decay of NMDA receptor-mediated synaptic currents [1]. The f-I curve is modeled by a smooth rectification function

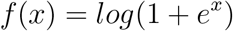

The bias term ***b_rec_*** admits per-unit firing thresholds. Intrinsic background noise current is modeled by an Ornstein-Uhlenbeck process ***ζ*** with time constant *τ_ζ_* and variance *σ_rec_*, where ***ξ*** represents the underlying independent white-noise process with zero mean and unit variance.

Output responses are readout from the activity of the RNN units by *N_out_* output units, ***y***, whose activity is given by

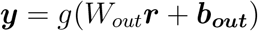

Here, *W_out_* is a *N_out_ × N* output weight matrix, ***b_out_*** is the bias of the output units, and 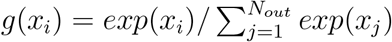 is the softmax or normalized-exponential function which produces output unit activity that indicates the probability of generating each of the *N_out_* response choices.

The model is simulated by temporal discretization of equation (1) with Euler’s method, as

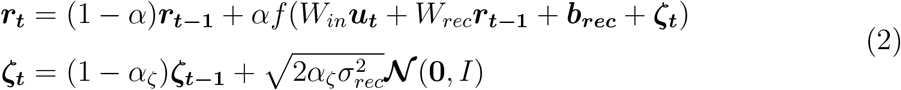

where the time-discretization step size is Δ*t*, *α* = Δ*t/τ*, *α_ζ_* = Δ*t/τ_ζ_* and ***𝒩*** (**0**, *I*) is a random vector sampled from a gaussian distribution with zero mean and identity covariance (*I*). In all figures, the network size *N* = 100, Δ*t* = 1 ms, *τ_ζ_* = 2 ms and *σ_rec_* = 0.05. The magnitude of the network- and input-unit firing rates is measured as the *L*^2^-norm of ***r_t_*** and ***u_t_***, respectively, and summarized by averaging over all time points in a trial.

### Task Structure (Fig. 1)

We trained the network model on a series of delayed sensorimotor association problems, one at a time. In each problem, the network had to learn a one-to-one correspondence between a pair of sample stimuli and a pair of motor responses. Each problem therefore comprised two trial types, one per stimulus-response pair. Each trial was 2 s in duration (*T* = 2), and started with a 500 ms sample epoch, followed by a 1 s delay epoch, and ended with a 500 ms choice epoch. During the sample epoch, the network concurrently received inputs representing a fixation stimulus and one sample stimulus. During the delay epoch, it continued to receive only the fixation input. It received no inputs during the choice epoch. The model was required to respond by maintaining fixation during the sample and delay epochs, and choosing the appropriate motor response during the choice epoch. Therefore, the model contained three output units (*N_out_* = 3), two to report response choices and one for fixation. This trial structure, including the available response choices, remained fixed across problems.

Sample stimuli were represented by ten-dimensional unit-length vectors (*L*^2^-norm = 1). The two sample stimulus input representations in a problem were drawn from a random gaussian distribution with zero mean and identity covariance. They were then orthogonalized to avoid learning efficiency confounds stemming from the relative difficulty in learning to distinguish between more versus less correlated sample stimuli. The fixation input was a scalar with value 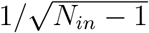 when it was on and zero when off. Therefore, there were a total of *Nin* = 11 input units. Learning-to-learn was robustly observed even in the absence of the orthogonalization step; however, the variance in learning efficiency was higher. Qualitatively similar learning-to-learn performance was also observed with 200-dimensional sample stimulus representations and *N* = 1000.

Each problem was learned over a sequence of trials, psueudorandomly sampled from the two trial types, until the average error on fifty consecutive trials fell below a criterion value (see *Network Training*). The learning efficiency for a problem was measured by the number of trials required to achieve this criterion. After a problem was learned, the model was transitioned to the next problem, wherein it had to learn to associate a new pair of pseudorandomly selected sample stimuli to the two motor responses.

### Network Training (Fig. 1)

A network was trained on a problem by updating its connection weights (*W_in_*, *W_rec_* and *W_out_*), biases (***b_rec_*** and ***b_out_***) and initial network state (***r*_0_**), so that it could choose the desired response for each of the sample stimuli. These updates were generated by stochastic gradient descent - an optimization algorithm that incrementally updates a network’s parameters at the end of each trial, based on the errors in the output unit responses during the trial. In contrast to standard RNN training practices wherein model parameters are adjusted based on the average error from a batch of several trials and learning efficiency is measured by the number of trial batches to reach criterion performance, our training procedure closely matched established animal training protocols and allowed learning efficiency to be measured by the number of trials to criterion performance. The backpropagation through time (BPTT) algorithm was used to resolve temporal contingencies while computing parameter updates. We additionally applied the ADAM optimizer [2] to enhance the efficacy of the updates. All networks were trained with a learning rate of 10^−4^, except in Supplementary Figure 1 where the learning rate was systematically varied. ADAM decay rates for the first and second moment estimates were set to 0.3 and 0.999, respectively, and the moment estimates were reset at the beginning of each problem. The model implementation and parameter update computations were performed with Tensorflow [3].

Prior to the first problem, a naive network’s input weights in *W_in_* were initialized with random values drawn from a gaussian distribution with zero mean and variance 1*/N_in_*; the recurrent weights in *W_rec_* were initialized with random values constrained by householder transformations such that the rows (and columns) of the initial recurrent weight matrix were orthogonal to each other and of unit length [4]. Initializing the recurrent weights in this manner allows gradients to be backpropagated more effectively. All other network parameters were initialized to zero. Upon transition to a new problem, all parameters retained their values. At initialization and throughout learning, the sign and sparsity of the weights and biases were not constrained. The initial network state was always restricted to non-negative values.

Network training was performed in a supervised setting, wherein the parameters were adjusted to minimize an objective function, *ℒ*, that included the errors in the model’s output responses:

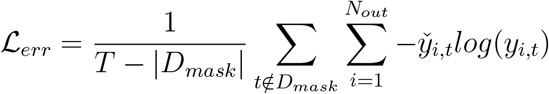

The error at each time step, *t*, was given by the cross-entropy of the probability distribution over responses generated by the network, ***y_t_***, relative to pre-specified target responses, ***y̌*_*t*_**. The total error for a trial, *ℒ_err_*, was the mean of the per-timestep error taken over the trial duration *T*. This mean excluded a masking interval, *D_mask_*, set to the first 100 ms of the choice epoch, which allowed for flexible reaction times. Networks were considered to have learned a problem when the average *L_err_* over fifty consecutive trials of the problem fell below a criterion value of 0.005.

The objective of the training procedure was to minimize the sum of this error and auxiliary regularization terms:

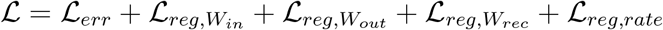

The regularization terms included both weight and activity regularization to encourage solutions that generalized well [5, 6] and generated stable network dynamics. We imposed *L*^2^ regularization on the input and output weights as follows:

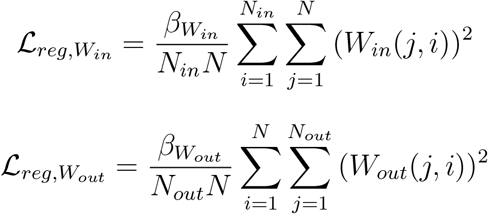

We observed that networks with a similar *L*^2^ regularization of the recurrent weights were sensitive to the value of meta-parameter *β_W_rec__*, particularly when the network size was large—small values of *β_W_rec__* produced unstable network dynamics during later problems, while large values hindered learning efficiency. The squared frobenius norm of the recurrent weight matrix, which constitutes such an *L*^2^ regularization, is given by:

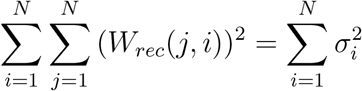

where *σ_i_* is the *i^th^* singular value of the recurrent weight matrix *W_rec_*.

An analysis of these singular values under conditions that led to unstable network dynamics revealed that their *L*^2^-norm (i.e. the square root of the right-hand side of the equation above) remained roughly fixed over the course of learning several problems; However, their distribution changed considerably across problems - smaller singular values shrank, while larger singular values grew and ultimately resulted in unstable network responses to novel sample stimuli. We mitigated this by introducing an alternate form of recurrent weight regularization that penalized the magnitude of the first *k* singular values of *W_rec_*:

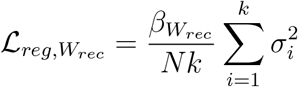

Finally, we imposed a homeostatic firing rate regularization:

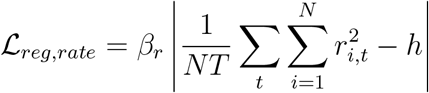

The meta-parameter *h* was set to zero for the first problem, effectively imposing an *L*^2^ regularization of the recurrent unit firing rates as the first problem was learned. To avoid unrestrained growth or reduction in the firing rates while learning subsequent problems, the homeostatic set-point *h* was then set to the mean squared firing rates averaged over the last fifty trials of the first problem. All networks were trained with *β_W_in__* = 10^−4^, *β_W_rec__* = 0.1, *β_W_out__* = 0.1, *k* = 10 and *β_r_* = 5 *×* 10^−4^, except in Supplementary Figure 1, where these hyper-parameters were systematically varied.

### Learning-to-learn Performance Characterization (Fig. 1)

A network exhibits learning-to-learn if its learning efficiency improves as a function of the number of previously learned problems. We evaluated this by quantifying the relationship between the trials-to-criterion on a problem and the number of problems learned thus far, where a decreasing relationship indicates learning-to-learn. Specifically, we fit a decaying exponential function to the number of trials to criterion *l*(*p*) on problem *p*, as a function of the number of learned problems *p −* 1:

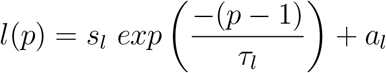

Here, *a_l_* represents asymptotic learning efficiency, *τ_l_* represents the time-constant to achieve this asymptote, and *s_l_*represents the improvement in learning efficiency between early and late problems. A large asymptote signifies poor learning-to-learn, while a large time-constant signifies slow learning-to-learn. The three parameters of the function were fit with the Levenberg-Marquardt algorithm implemented by the fit function of MATLAB’s curve fitting toolbox. The learning efficiency on the first problem was excluded from this analysis.

### Subspace Decomposition (Fig. 2)

We performed semi-supervised dimensionality reduction on the population activity, to determine how strongly and consistently the shared task structure is represented across problems. The procedure begins by compiling a tensor *R_k,t,j,i_* of activity patterns generated by the population of firing rate units (*k* ∈ [1, *N*]) over time (*t* ∈ (0, *T*]), for the two *response* types (*j* ∈ {*response*_1_, *response*_2_}) across a group of fifty consecutively learned problems (*i ∈* [*p* + 1, *p* + 50]). This assembles a tensor of one hundred population trajectories for the group, fifty for each response type. The semi-supervised dimensionality reduction extracts decision representations that are shared by the group as follows. Stimulus- and problem-specific representations for each response type are averaged out, or marginalized, across problems in the group:

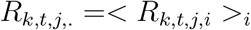

Principal components analysis is performed on a concatenation of the resulting two trajectories in *R_k,t,j,._*. The loading vectors for the first *m* principal components are collected into a *N × m* loading matrix *L_D_*. These vectors define a basis for the decision subspace. Importantly, to ensure that the decision subspace fully captures shared decision representations, the marginalized trajectories are not de-meaned before performing principal components analysis. Here, we set *m* to 4, as the first 4 principal components collectively explained at least 98% of the variance in the marginalized trajectories, in all the networks we analyzed.

Next, an *N × N* projection matrix *P* (*Q*) that projects population activity into the decision subspace (stimulus subspace), is defined as:

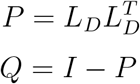

where *I* is the identity matrix. The decision components of the learned trajectories for problem *p* + *x* (*x ∈* [1, 50]) are identified as:

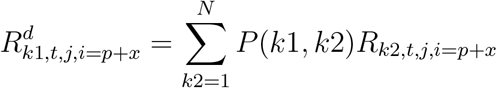

and their stimulus components as:

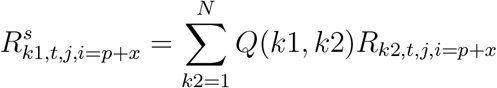

where *P* (*k*1, *k*2) and *Q*(*k*1, *k*2) represent the element in the *k*1*^th^* row and *k*2*^th^* column of the respective projection matrices. The decision components are further decomposed into mean 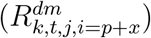 and residual 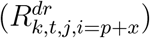 decision components, as:

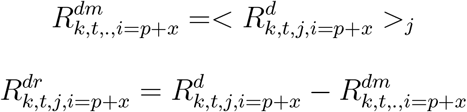

The net current from these components 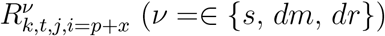 to an output unit *o* was computed as 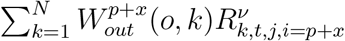 where 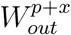 is the output weight matrix learned in problem *p* + *x*. The dimensionality of any set of vectors (e.g. population activity in the stimulus subspace) was approximated by its participation ratio [7], computed as 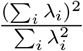, where *λ_i_* is the *i^th^* eigenvalue of the covariance matrix of the vectors.

### Manifold Perturbations (Fig. 3)

To assess whether the reuse of the decision representations improves learning efficiency, networks were trained on their second problem while constraining them in a manner that required the formation of new decision representations. The learning efficiency of such networks was compared to controls that were allowed to reuse existing decision representations while learning their second problem.

A naive network was first trained on 50 problems and the corresponding populations trajectories were used to identify its decision and stimulus subspaces. The network’s parameters, including its output weights, were reset to their values at the end of the first problem. Then, its output weights were perturbed, and the network was trained on a new problem, i.e. a second problem with respect to its parameters, while barring the training procedure from changing its output weights. This procedure was repeated fifty times for each network, resetting its parameters, applying an independently chosen random perturbation to its output weights, freezing the output weights, and training the network on a new sample stimulus pair each time. The output weights were subjected to one of three forms of perturbation. In the frozen readout condition, the output weights were unperturbed after the parameter reset. In D *→* S manifold perturbations, following the parameter reset, the output weights were perturbed to replace the overlap between the network’s readout and decision subspaces with a corresponding overlap between its readout and stimulus subspaces:

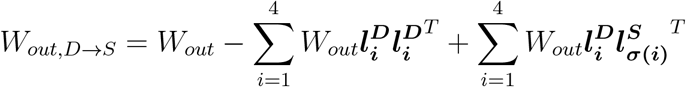

where *W_out,D→S_* is the perturbed output weight matrix, 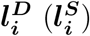 is the *i^th^* principal component loading vector of the decision (stimulus) subspace, and *σ*() represents a random shuffle or permutation of the stimulus subspace principal component loading vectors. In S *→* S manifold perturbations, following the parameter reset, the output weights were perturbed to permute the overlap between the readout and stimulus subspaces:

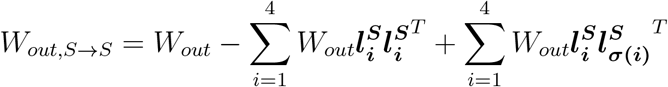

### Relationship between pre-learning and learned trajectories based on weight- and state-driven vector field changes (Figs. 4 - 5)

Over the course of learning problem *p*, the model’s parameters change from their values at the beginning of the problem, i.e. their pre-learning values 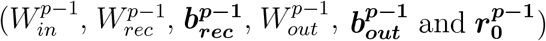, to their values at the end of the problem, i.e. their learned values 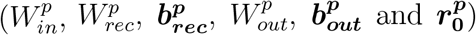. The difference between the learned and pre-learning values of the parameters quantify their change due to learning problem 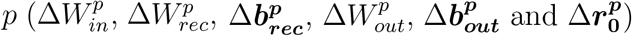, and are collectively referred to as Δ*W^p^*.

Due to these parameter changes, the population activity in response to inputs 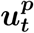 is altered from its pre-learning levels, 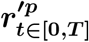, to its learned ones, 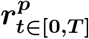 (Fig. 4c, left). We derive an expression for this change in population activity, 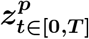, in terms of the parameter changes. Based on the time-discretized model equation (2), we have:

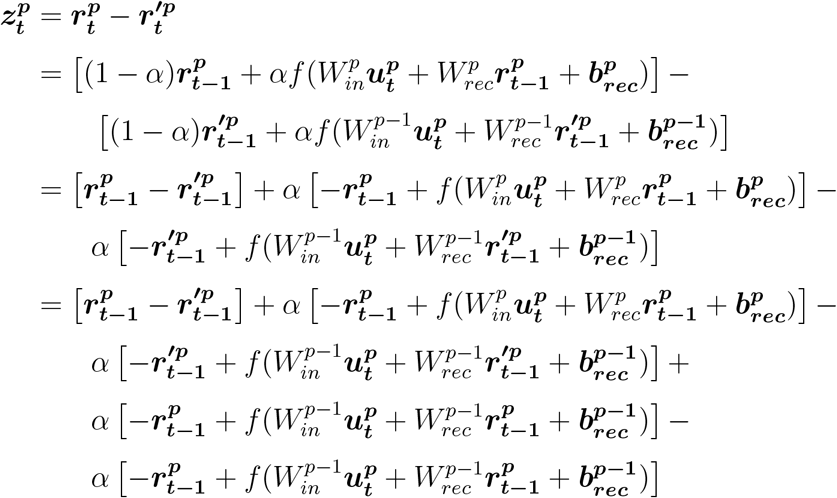

Rearranging the terms, we have:

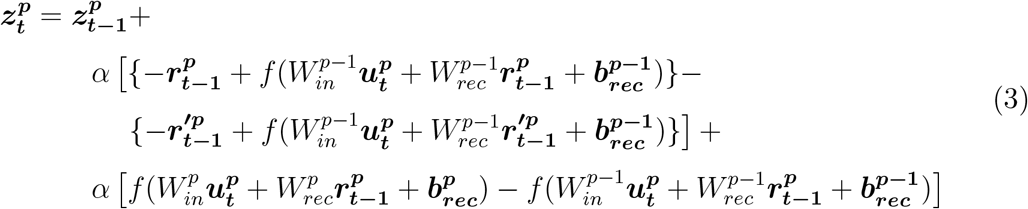

This expression shows that the change in population activity emerges from an accumulation of activity change increments, 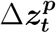 (Fig. 4c, center):

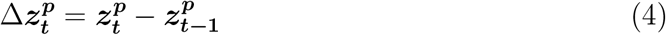

These increments are composed of two terms:

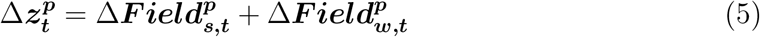

The first term, 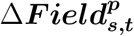, expresses the difference in the pre-learning vector field at the positions in state space along the learned 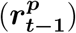 and pre-learning 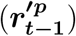 trajectories (Fig. 4d, left). It is therefore referred to as the *state-driven vector field change* (or state-driven VFC):

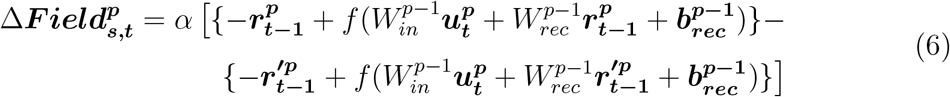

The second term, 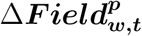, expresses the change in the vector field at population states along the learned trajectory due to the parameter changes (Fig. 4d, center; Fig. 4b, right). It is therefore referred to as the *weight-driven vector field change* (or weight-driven VFC):

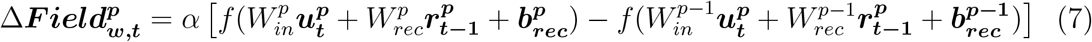

The weight-driven VFC stems from the change in the net afferent currents to the population, 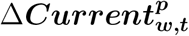, due to the parameter changes (Fig. 4b, left):

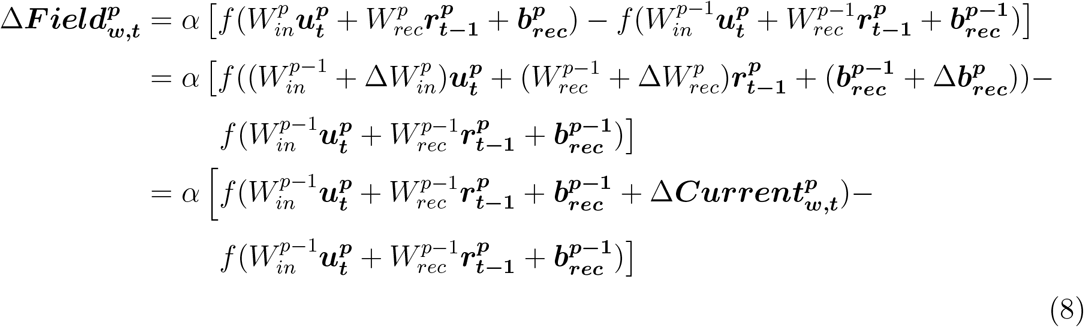

where 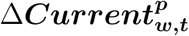 is determined by 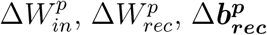, as:

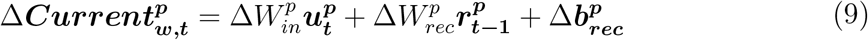

The change in initial population state is defined as 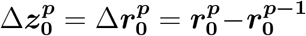. We omit the contribution of this change from our analyses, as it consistently showed a negligible effect on the evolution of the learned trajectory and the activity changes, across all problems and networks tested.

The contribution of the two vector field change terms to the activity change increment, 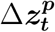, was measured by their magnitude along, or in the direction of, 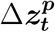 (Fig. 5a). This was computed by vector projection, as:

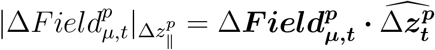

where *µ ∈* {*w, s*}, ***·*** represents the dot product operator, and 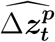 is the unit vector in the direction of 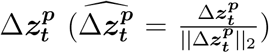. Therefore, the vector field change along 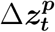 is given by:

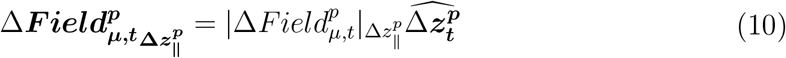

The remainder of each vector field change term represents its components orthogonal to 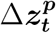 (Fig. 5a):

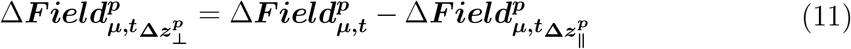

In order to compare the relative direction of the orthogonal components of the weight- and state-driven VFCs (Fig. 5a), we arbitrarily (but without loss of generality) chose the direction of 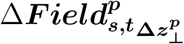 as the reference — signed magnitudes were computed by vector projection of 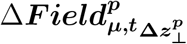 onto a unit vector in the direction of 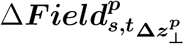.

The magnitude of change in the input and recurrent connection weights was measured by their frobenius norm, 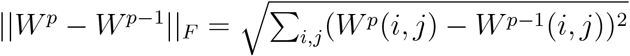.

### Contribution of changes in individual parameters to the weight-driven vector field change (Fig. 6)

We measured the individual contributions of changes in the input weights 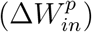, recurrent weights 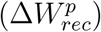 and network unit biases 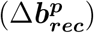 to the weight-driven vector field change. Note that, the postsynaptic current changes can be linearly decomposed based on the contributions of these parameter changes: they are given by the 3 terms on the right-hand side of equation (9), which we denote as 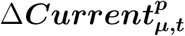 to signify postsynaptic current changes due to changes in the parameter 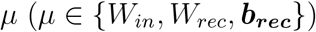. In contrast, the vector field change is a non-linear function of these parameter changes. Therefore, we formulated non-linear approximations of their contributions, which we denote as 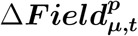. Such an approximation must solely depend on changes in the parameter *µ*, and the approximations must collectively satisfy the following with an acceptably small approximation error:

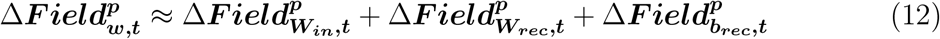

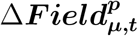 is a vector, whose elements 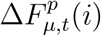, represent the approximate change in the firing rate of network unit *i*, at time *t* along the learned trajectory for problem *p*, due to the changes in parameter *µ*. From equations (8) and (9), we observe that it is related to the net postsynaptic current at unit *i* due to the pre-learning parameter values, 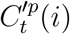, and the net change in the postsynaptic current at unit *i*, 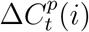. These net currents are given by the summations 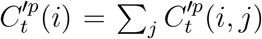 and 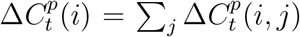, where *j* denotes an individual connection/bias that contributes to the net current into unit *i*. These include its individual afferent input and recurrent connections weights and its bias. We use the notation 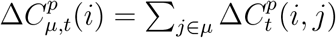 to explicitly refer to the net contribution of changes in parameter *µ* to the postsynaptic current changes at unit *i*. We derive an expression for 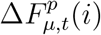 via taylor-expansion of equation (7):

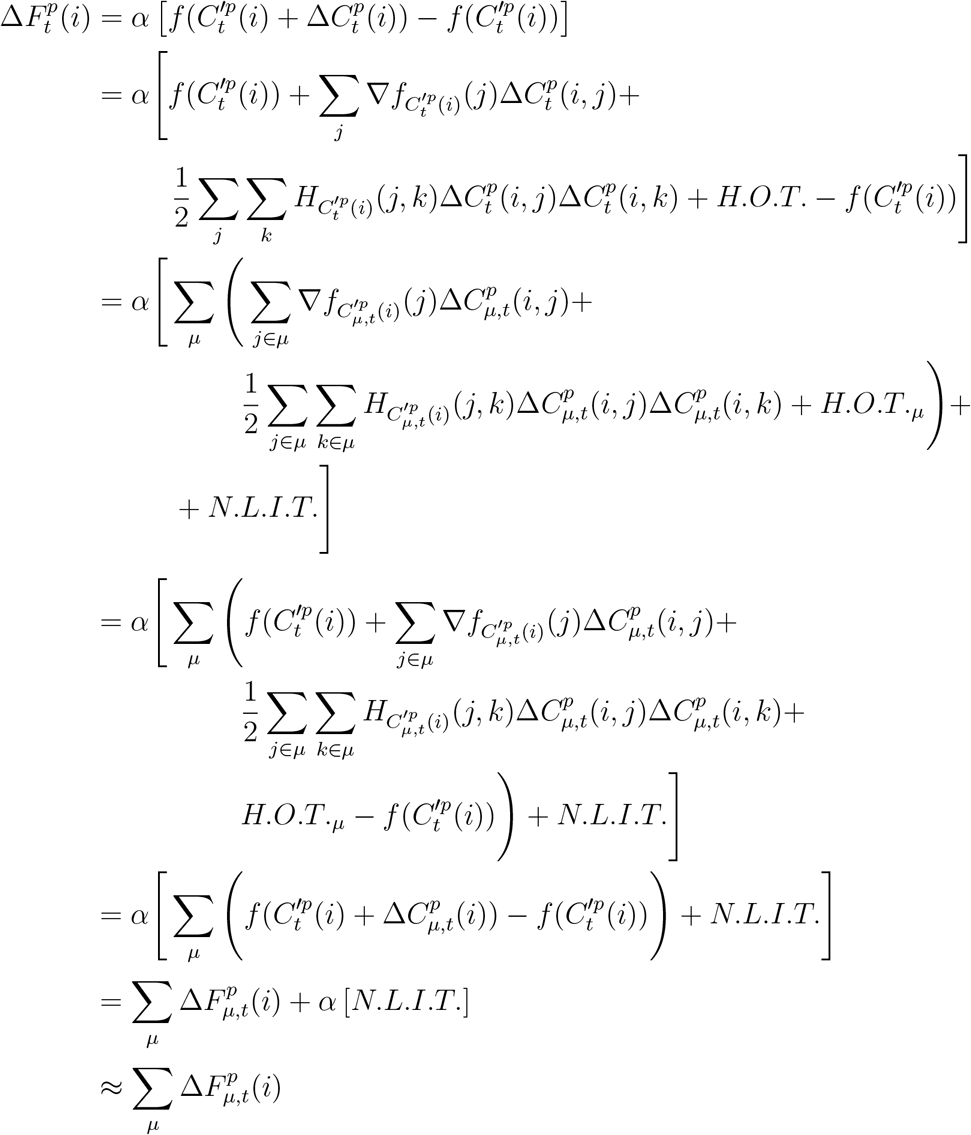

where 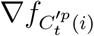 and 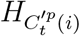 correspond to the gradient and hessian of *f* for unit *i*, when the magnitude of its net postsynaptic current is 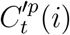. *H.O.T.* corresponds to the higher-order terms of the taylor expansion, *H.O.T._µ_* corresponds to the higher-order terms that only involve changes to parameter *µ*, and *N.L.I.T.* corresponds to non-linear interactions between the terms due to changes in *W_in_*, *W_rec_* and ***b_rec_***. From the derivation above, we have:

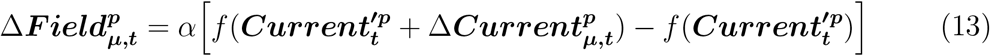

This equation expresses the unique contribution of changes in the parameter *µ* to the vector field change. Furthermore, the collective contribution of the changes in the three parameters satisfy equation (12), subject to an approximation error of *α**N.L.I.T.***. We calculated the magnitude (*L*^2^-norm) of this error; at each trial timestep and in each network tested, this error was found to be less than 1% (average across problems). In supplementary figures 5 - 6, we forego presenting the contribution of the change in network unit biases 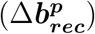, as it consistently showed a negligible effect on the changes in postsynaptic currents and the vector field in all problems and networks tested.

For figure 6, this approach was extended to estimate the unique contribution of the changes in recurrent connection weights from the decision 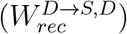 and stimulus 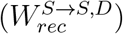 subspaces to the vector field change, where 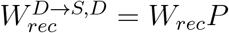 and 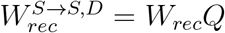. To do so, we extended the parameter set in the derivation above to 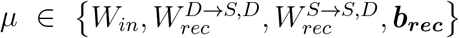. Also, the decision and stimulus components of the vector field change due to recurrent weight changes were calculated as 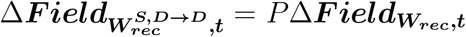 and 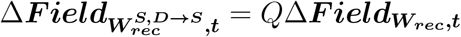, respectively.

The clamping simulations to evaluate reciprocal interactions between stimulus and decision representations (Supplementary Fig. 8c) were performed as follows. Starting from the initial population state (***r*_0_**), the model was simulated for a single timestep with the learned parameter values as per equation (2). This advanced the population state to ***r*_1_**. The stimulus (decision) representation was then reset to its pre-learning value 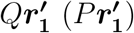, and the model were simulated for another timestep. This process was repeated until the end of the trial. The Euclidean distance (or magnitude of deviation) between the decision (stimulus) representations observed during these simulations and the learned decision (stimulus) representations reflected the strength of the reciprocal interactions.

### Relationship between the accumulation of weight changes across problems and the progressive decrease in the weight-driven vector field change (Fig. 7)

We measured the contribution of the weight changes elicited while learning problem *p−k* (Δ*W^p−k^*, for 1 *≤ k ≤ p −* 2) to the cumulative vector field change (or cumulative VFC) along the learned trajectory for problem 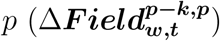 as:

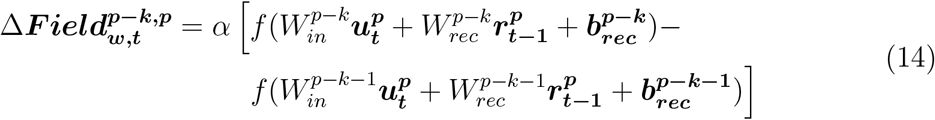

Then, the cumulative vector field change due to the accumulation of weight changes across all the learned problems from *p − k* to *p −* 1 was given by:

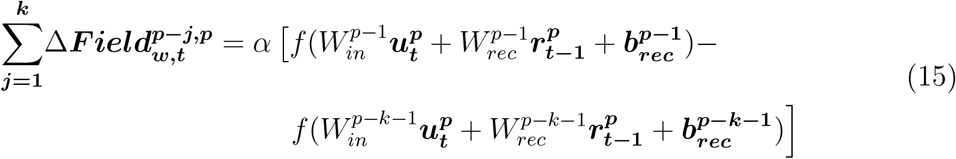

In figure 7, we present the magnitude of cumulative VFC along the parallel 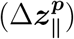 and orthogonal 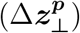 components of the vector field change for problem *p*. These were computed via vector projection of the cumulative VFC onto unit vectors in the direction of the vector field change components. Specifically, given that the vectors 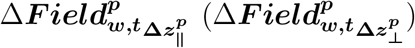 are nearly one-dimensional across trial time *t* within problem *p* (Fig. 5c), we applied principal components analysis to find a single basis (unit-norm) vector, 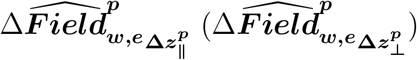, that accurately represents their shared direction during each non-overlapping 250 ms epoch, *e*, of the trial. The magnitude of the cumulative change along the parallel / orthogonal vector field change component was given by:

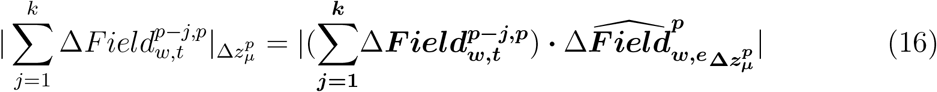

where *µ ∈* {∥, ⊥}, and time *t* lies within the interval of epoch *e*. The magnitudes of cumulative VFC contribution by individual problems along the parallel / orthogonal vector field change component 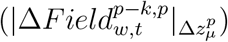 were computed similarly.

The signed cumulative VFC and per-problem cumulative VFC contributions in figure 7c were calculated as above, but without taking the absolute value on the right-hand-side.

The per-trial magnitude of the cumulative VFC contribution by problem *p − k* to problem *p* was calculated as 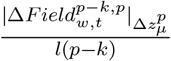, where *l*(*p − k*) is the trials-to-criterion for problem *p − k*. The sum of the magnitudes of the cumulative VFC contributions to problem *p* was calculated as 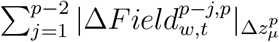.

The magnitude of net suppression of problem *p*’s weight-driven VFC along its parallel / orthogonal component is defined as the net suppression in the direction of the corresponding component due to net weight changes between the start of problems 2 and *p*. It was computed from the total vector field change along the learned trajectory for problem *p* since the start of problem 2. Let 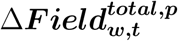 represent this total vector field change at time *t*:

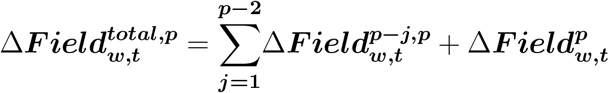

Then the total change along the parallel / orthogonal vector field change component was given by:

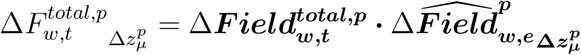

We applied a sign correction to this quantity to ensure that its temporal mean is always positive. This allowed us to accurately calculate the net suppression. After sign correction, 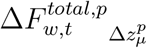 becomes:

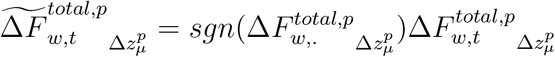

where 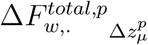 represents the temporal mean of 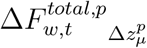 over time *t* within a trial, and *sgn*() represents the signum function. Similarly, the weight-driven VFC for problem *p* along its parallel / orthogonal components was given by:

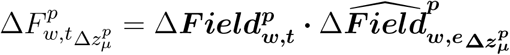

Then, the magnitude of net suppression along the parallel / orthogonal vector field change component for problem *p* was:

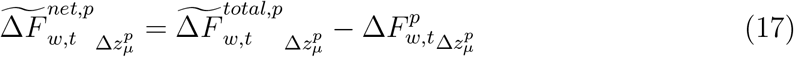

The progression of this quantity over the learning-to-learn timecourse can be described in terms of the number of previously learned problems. We note that the temporal mean of the magnitude of the weight-driven VFC along its parallel / orthogonal component 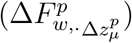 decays exponentially from problem 2 onwards until an asymptotic value *b_µ_* is converged upon (as in Fig. 6b). This decay may be expressed as:

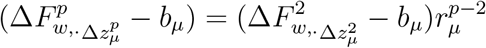

for an appropriate base *r_µ_ <* 1. Taking the temporal mean of equation (17) over trial time *t*, we have:

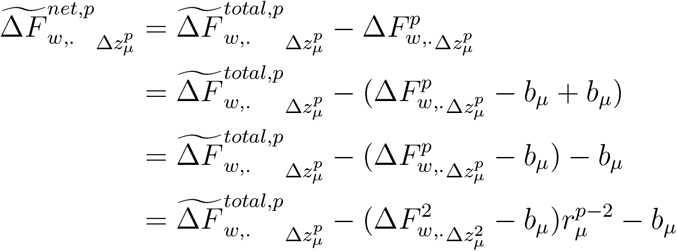

Rearranging, we have:

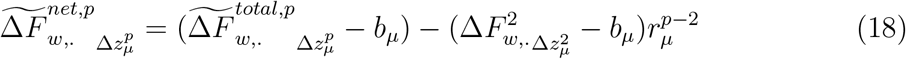

This equation expresses the progression of the magnitude of net suppression over the learning-to-learn timecourse, and determines its shape as a function of the number of previously learned problems (Fig. 7b). Note that when the first term 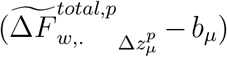 is roughly constant across learning-to-learn stages (as we found by measurement), the magnitude of net suppression is given by an inverted exponential function.

Finally, we determined the relative contributions of the cumulative input versus recurrent weight changes to the cumulative VFC along the orthogonal vector field change component (Supplementary Fig. 9c). To do so, we calculated the cumulative VFC for problem *p* solely due to the accumulation of input weight changes elicited by previously learned problems as:

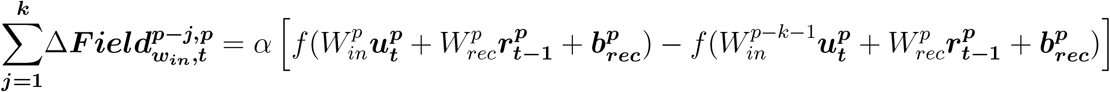

The cumulative VFC solely to due to recurrent weight changes was calculated similarly. Both quantities were then projected onto the basis vector for the orthogonal vector field change components in problem *p* (as in equation (16)), to compare their contributions along this component.

## Supplementary Notes

### 1.1 Recurrent plasticity elicits efficient learning, but is not necessary for it

The reuse of the decision manifold to compose learned trajectories implies that a problem is learned by mapping each of its sample stimuli onto an appropriate decision representation within the decision manifold. The model may achieve this either by adjusting its input connection weights to appropriately remap the novel sensory inputs, or by adjusting its recurrent connection weights to alter how these inputs are recurrently integrated into the appropriate decision representations, or by some combination of the two. To assess the relative contributions of these two mechanisms, we compared: (i) the magnitude of change in the input and recurrent weights when problems are learned; (ii) the decrease in output accuracy when the input or recurrent weights changes are reversed; (iii) the learning-to-learn performance of networks with a pre-established representational manifold, that must exclusively rely on changes to either their input or recurrent weights to learn new problems.

We observed that the input weight changes were similar in magnitude to the recurrent weight changes (Supplementary Fig. 5a). Yet, reversing these relatively large input weight changes produced a negligible decrease in response accuracy. In contrast, reversing the recurrent weight changes decreased response accuracies to chance levels (Supplementary Fig. 5b). To address this discrepancy between the relative magnitude of the weight changes and their effect on output response accuracy, we approximated and compared the individual contributions of the input and recurrent weight changes to the weight-driven VFC (equation (13), see *Methods*). Consistent with the latter result, we found that the weight-driven VFC is primarily caused by recurrent weight changes (Supplementary Fig. 5c, left). Recall that the weight-driven VFC is directly related to changes in the postsynaptic currents (Fig. 4b, equation (8)), which is a product of the connection weight changes and the firing rates of network and input units (equation (9)). Given that the input and recurrent weight changes are comparable in magnitude, we posited that the disproportionate contribution of the recurrent weight changes would be explained by a difference in the magnitudes of the network and input unit firing rates. A comparison of these firing rate magnitudes confirmed our hypothesis (Supplementary Fig. 6d, Input Act. Mag. = 1, *W_in_* gain = 1, Baseline).

These results demonstrate that the model prefers solutions that rely on recurrent weight changes, and that these solutions make more efficient use of the weight changes to alter the vector field. But are these solutions preferred because they are more efficient to learn? In other words, do solutions that rely on input weight changes exhibit poor learning efficiency? To answer this question, we trained networks with pre-established decision and stimulus manifolds (i.e. networks trained on their first problem) on new problems, either with frozen input weights or with frozen recurrent weights. We then compared their asymptotic learning efficiency. Indeed, we found that networks with frozen recurrent weights exhibited substantially higher learning efficiency asymptotes than networks with frozen input weights (Supplementary Fig. 6c)—the model’s preference for solutions that relied on recurrent weight changes was predicated on their superior learning efficiency. Moreover, networks with frozen recurrent weights required considerable changes to their input weights before they had learned a problem (Supplementary Fig. 6c, Input Act. Mag. = 1, *W_in_* gain = 1). This suggests that the model’s learning efficiency on a problem is related to the magnitude of connection weights changes that are necessary to learn it. We further explore this relationship in the main text (Fig. 6).

In the networks explored thus far, learning is more efficient when it relies on recurrent weight changes. We sought to understand whether this is always true, i.e. are recurrent weight changes a necessary condition for efficient learning? Or, do network regimes exist wherein learning is equally efficient when driven by input weight changes? Networks that learn via input weight changes exhibit poorer learning efficiency due to deficits in their influence on the postsynaptic current changes and the weight-driven VFC (Fig. 4b). Therefore, we reasoned that such networks may become efficient learners in regimes where this deficit is eliminated. To test this, we measured the asymptotic learning efficiency of networks with a tenfold increase in input unit firing rates, and with frozen recurrent weights. We expected that this increase would facilitate a stronger influence of input weight changes on postsynaptic current changes, thereby improving learning efficiency. Surprisingly, we found that these networks also under-performed in comparison to networks with plastic recurrent weights (Supplementary Fig. 6b, Input Act. Mag. = 10, *W_in_* gain = 1). Consistent with the original networks (Input Act. Mag. = 1, *W_in_* gain = 1), they required considerable input weights changes (Supplementary Fig. 6c) to generate similarly-sized postsynaptic current changes as fully plastic networks (Supplementary Fig. 6e). Again, this was due to an asymmetry in the input and network unit firing rate magnitudes (Supplementary Fig. 6d): the network unit firing rates further increased in response to the elevated input unit firing rates, because the network units were receiving larger currents from the input units.

Finally, we reasoned that this increase in the network unit firing rate magnitudes could be avoided by additionally scaling down the magnitude of the input weights. This would both generate similarly-sized efferent currents from the input units as in our original networks, and facilitate a stronger influence of input weight changes on the postsynaptic current changes due to the elevated input unit firing rates. We tested this in networks with both a tenfold increase in the input unit firing rates and a 20-fold decrease in the initial input weights (i.e. input weights of the naive network). We now found that networks with frozen recurrent weights exhibited learning efficiency asymptotes that were comparable to their fully plastic counterparts (Supplementary Fig. 6b, Input Act. Mag. = 10, *W_in_* gain = 0.05). These networks produced input weight changes of comparable magnitude to the weights changes in fully plastic networks (Supplementary Fig. 6c), while also producing postsynaptic current changes of comparable magnitude (Supplementary Fig. 6e). This was because input and network unit firing rates were comparable in magnitude (Supplementary Fig. 6d). Taken together, these analyses demonstrate that fully plastic networks learn new problems largely via recurrent connection weight changes because it is generally more efficient to do so. However, recurrent weight changes are not necessary for efficient learning: Network regimes exist wherein learning via input weight changes is equally efficient.

### 1.2 Plasticity alters stimulus representations far more than decision representations

We have demonstrated that the model learns a problem when each sample stimulus elicits decision and choice representations that appropriately direct the desired output response, and that this is achieved by the preferential engagement of plasticity in the network’s recurrent connections. But does this process also enlist and alter the stimulus representations, and if so, to what end? Measurements showed that both the decision and the stimulus representations developed sizeable changes after learning (Supplementary Fig. 8b). To test the utility of the changes in the stimulus representations, we simulated trials in a network that had learned a problem, while clamping its stimulus representations at their pre-learning values (see *Methods*), and measuring the effect of this intervention on the decision representations. If the learned decision representations evolve independently of the stimulus representations, they should remain largely unaltered. Instead, we found that the decision representations experienced large deviations (Supplementary Fig. 8c). Similarly, clamping the decision representations at their pre-learning values produced large deviations in the stimulus representations. This shows that the stimulus and decision representations sustain strongly recurrent interactions, and that changes in the stimulus representations are necessary both to remap sensory inputs onto the appropriate decision manifold and to maintain these decision representations throughout the trial.

We also examined whether the decision and stimulus representations mutually influence each other’s weight-driven VFC. Specifically, how is the weight-driven VFC modulated by pre-synaptic population activity in the stimulus versus decision subspaces? And to what extent does the resulting weight-driven VFC alter subsequent stimulus versus decision representations? In figure 6c, we show that the weight-driven VFC is primarily modulated by pre-synaptic population activity in the decision subspace, i.e. the decision representations predominantly scaffold the weight-driven VFC. Moreover, this decision- and weight-driven vector field change is primarily responsible for learning — reversing it reduced output accuracy almost to chance levels, while reversing the stimulus- and weight-driven vector field change had a much weaker effect (Supplementary Fig. 8d). Finally, a comparison of the overlap of the weight-driven VFC with the stimulus versus decision subspaces showed that weight changes mostly alter stimulus representations (Supplementary Fig. 8e).

These results suggest that reciprocal interactions between the stimulus and decision representations play a key role not only in decision making and working memory maintenance of these decisions, but also in learning the two mappings in each problem. They further demonstrate that the decision representations scaffolds the weight-driven VFC, and that the weight-driven VFC largely prevents uncharacteristic changes to the existing stimulus representations, a finding that is consistent with our results in figure 5 and supplementary figure 9a.

## Supplementary Figures

**Supplementary Figure 1:**
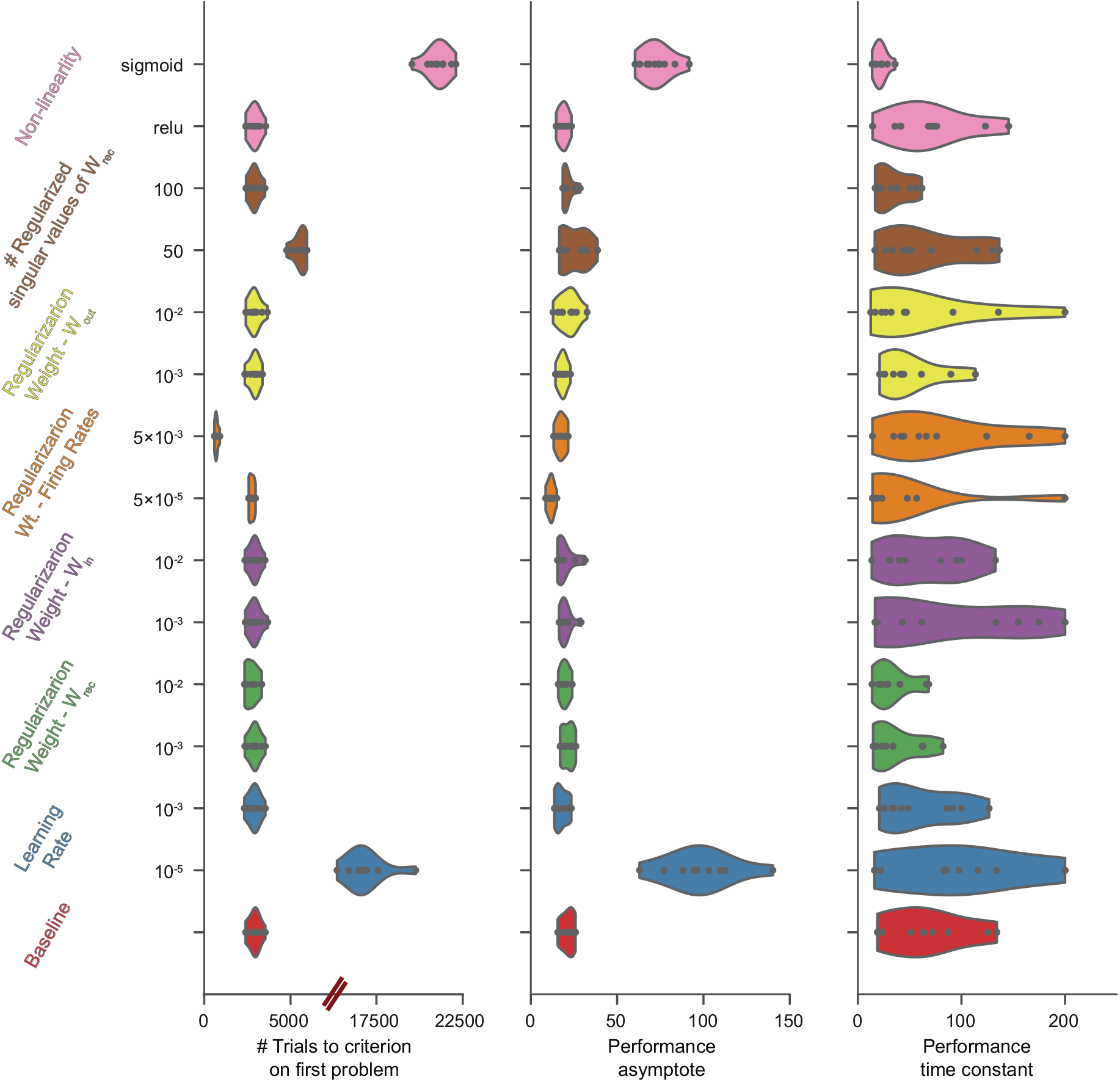
Learning-to-learn is robustly observed across a range of hyper-parmeter settings. Problem 1 learning efficiency (left), learning efficiency asymptotes (middle), and learning efficiency time constants (right), of networks trained with different learning rates, recurrent (β_Wrec_), input (β_Win_) and output (β_Wout_) weight regularization levels, numbers of recurrent weight matrix singular values (k) that are regularized, firing rate regularization levels (β_r_), and f-I transfer functions. Performance measures are presented in comparison to the baseline networks discussed in the main text. The regularization hyper-parameters and learning rates spanned 2 orders of magnitude. 10 networks were trained per hyper-parameter setting. Networks with sigmoid f-I transfer functions were trained with input unit firing rates scaled up by a factor of 10. Networks at the slowest learning rate and those with a sigmoid f-I transfer functions exhibited slower learning. However, all networks demonstrated learning-to-learn.

**Supplementary Figure 2:**
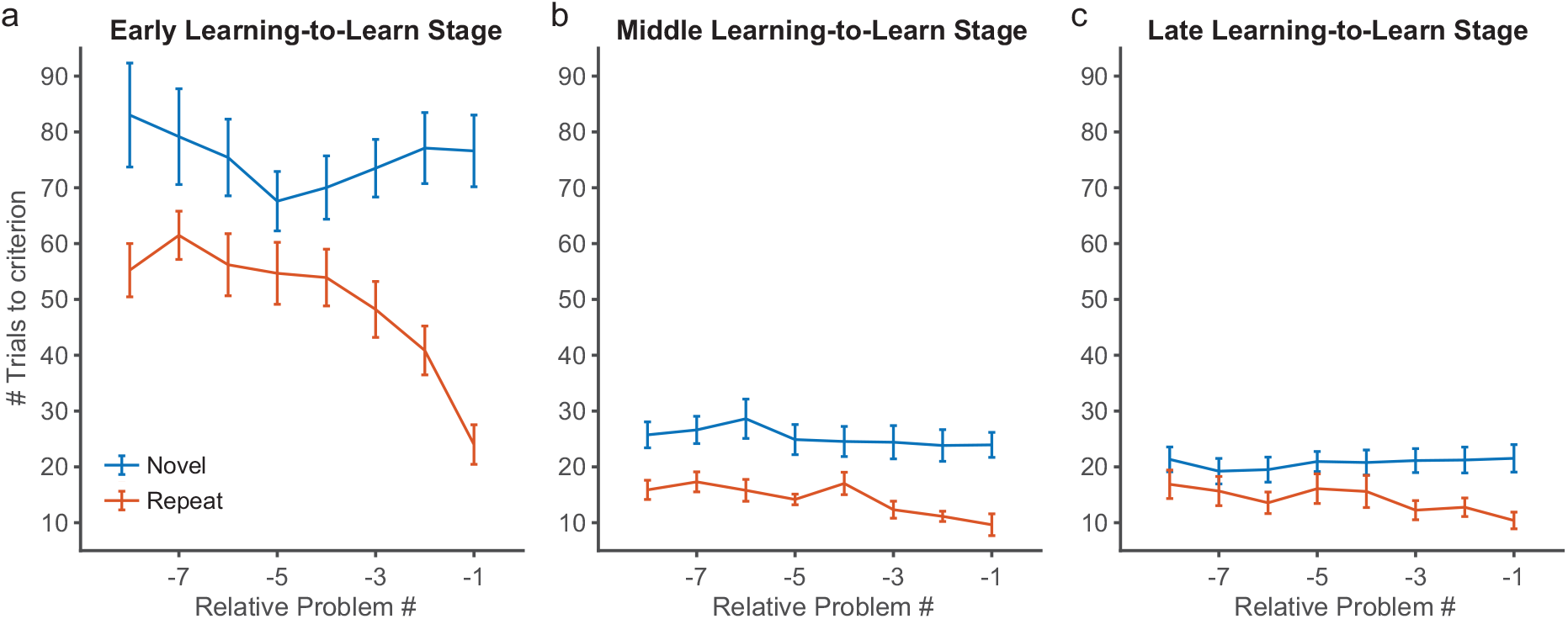
Model retains a memory of past problems. **a.** Learning efficiency of new problems (novel condition, blue) at early-stage learning-to-learn (20 problems in the range 11 - 55), in comparison to the learning efficiency when re-learning them following a varying number of intervening problems (repeat condition, red). **b-c.** Similar comparison of novel versus repeat learning efficiency for problems at middle-stage (**b**, 20 problems in the range 146 - 205) and late-stage (**c**, 20 problems in the range 346 - 405) learning-to-learn. Plots reflect mean values over 10 networks with different initial conditions, and error bars indicate the standard error of the mean.

**Supplementary Figure 3:**
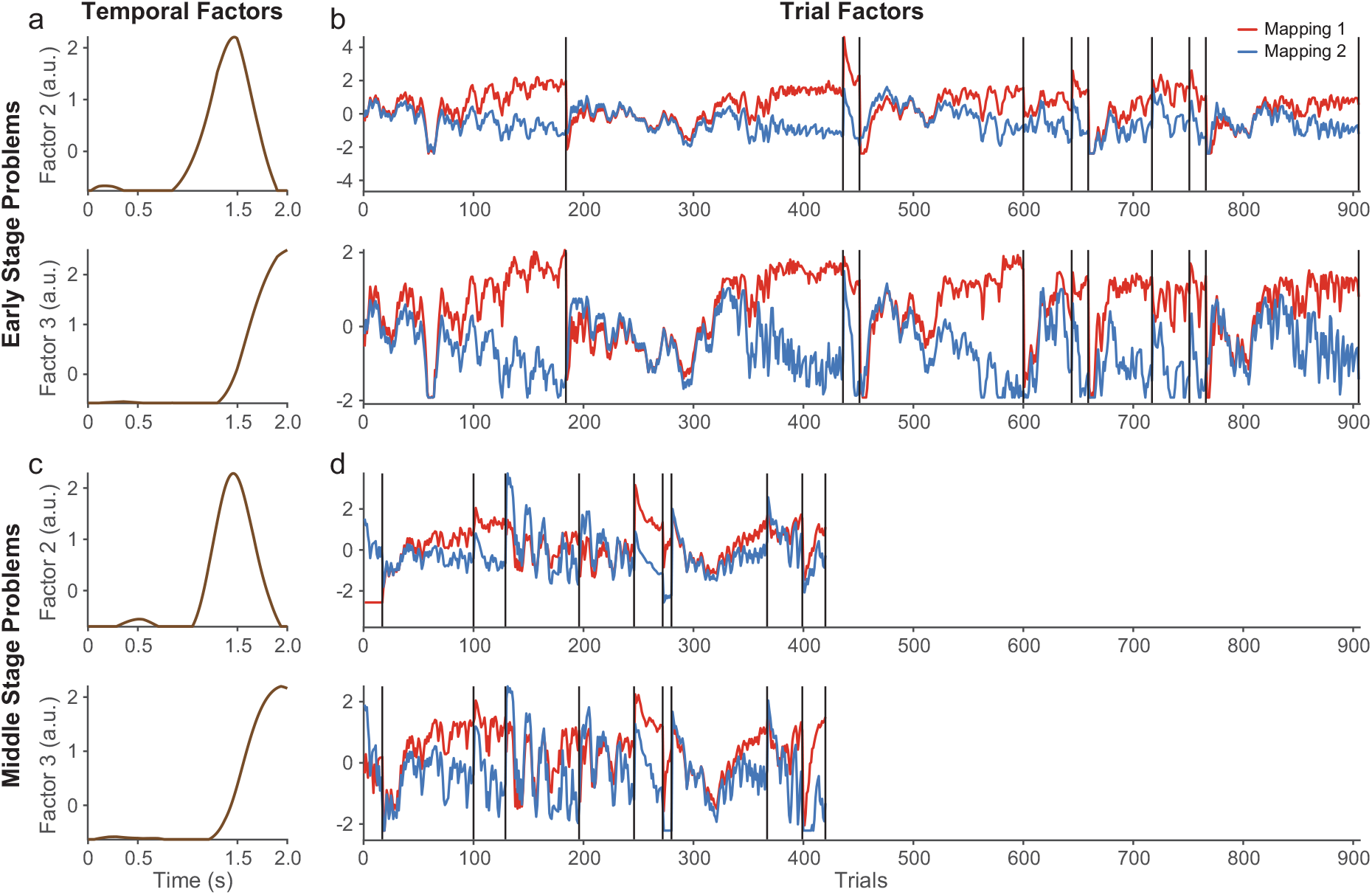
Learning dynamics are largely comprised of changes in shared population representations across problems. **a-b.** Temporal (**a**) and trial (**b**) factors produced by tensor decomposition analysis ([1]) when applied to population activity during trials between the beginning and end of learning 10 consecutive problems at early-stage learning-to-learn (problems 2 - 11). **c-d.** Temporal (**c**) and trial (**d**) factors of population activity during learning trials of problems at middle-stage learning-to-learn (problems 96 - 105). Plots reveal the emergence of large changes in delay- and choice-epoch activity as the problems are learned. These changes separate the population activity for the two mappings in a problem, in a manner that is consistent across problems. In addition, they emerge more rapidly while learning problems at middle-stage learning-to-learn.

**Supplementary Figure 4:**
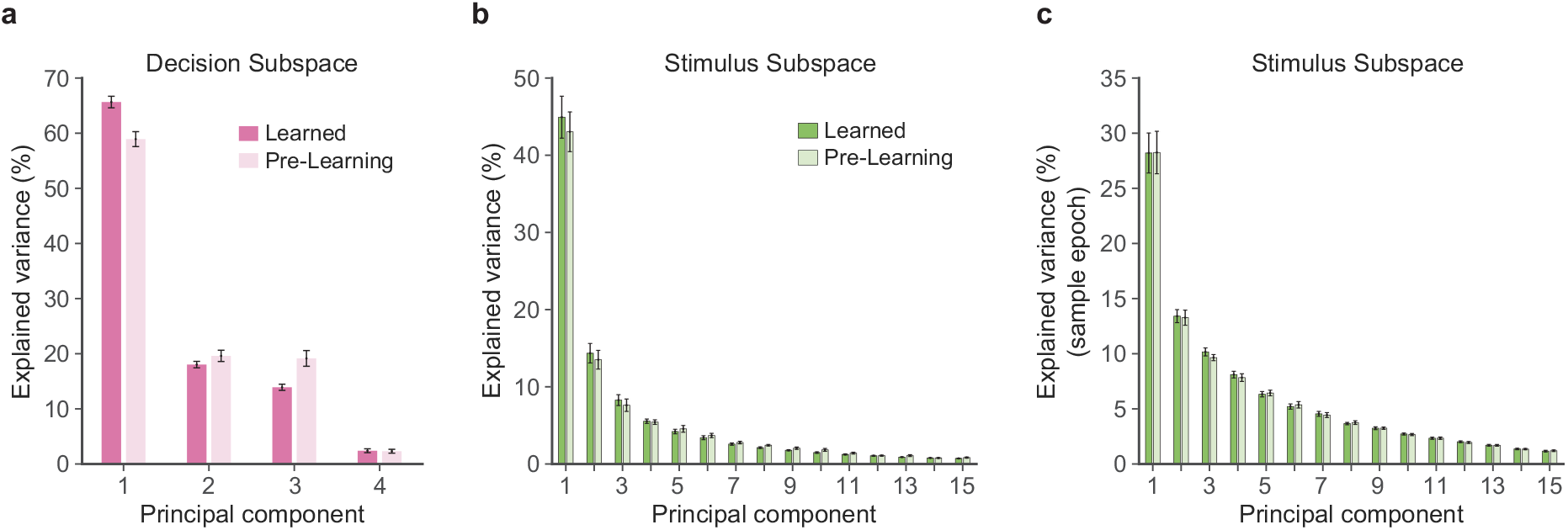
Intrinsic structure of the learned neural representations is also recruited in response to novel sample stimuli. **a.** Comparison of the variance in pre-learning and learned population activity that is explained by each principal component of the decision subspace. The decision subspace and its principal components were computed from the learned population activity across 50 consecutive problems. The variance in both the pre-learning and learned activity was measured along these principal components. **b-c.** Comparison of the variance in pre-learning and learned population activity that is explained by the first 15 principal component of the stimulus subspace, measured across the entire trial duration (**b**) and across the sample epoch of the trials only (**c**). The structure underlying learned neural representations is recruited even at the start of each problem, when the sample stimuli presented to the network are novel. The first problem was excluded from the pre-learning variance measurements, as the decision and stimulus representations develop only after the first problem is learned. Bars show mean variance explained are across 10 networks with different initial conditions, and error bars indicate standard errors of the mean.

**Supplementary Figure 5:**
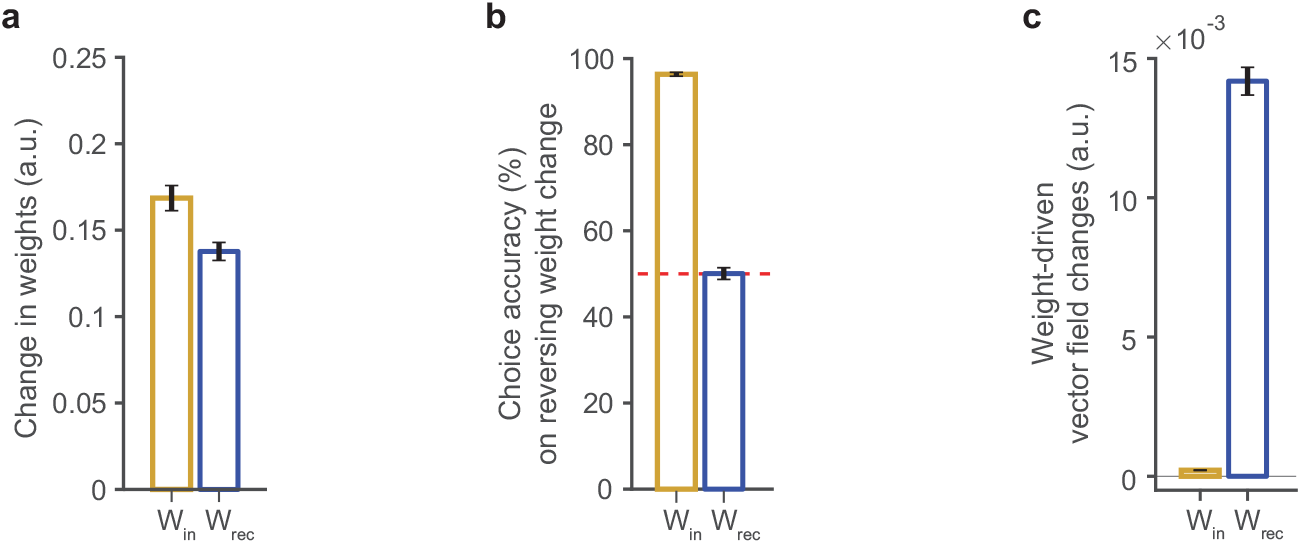
Learning largely relies on changes in the recurrent connection weights. **a.** Magnitude of the recurrent and input connection weight changes, measured by their Frobenius norms. **b.** Output accuracy when either the input or recurrent connection weight changes are reversed. **c.** Approximate magnitude of the weight-driven vector field change due recurrent and input connection weight changes. Magnitudes shown are the temporal mean of the L^2^-norm of the corresponding vector quantities over the entire trial duration, and averaged over both mappings in a problem. All measures presented are averages across problems 2 thru 51. All bars represent the mean across 10 networks with different initial conditions, and error bars indicate their standard errors.

**Supplementary Figure 6:**
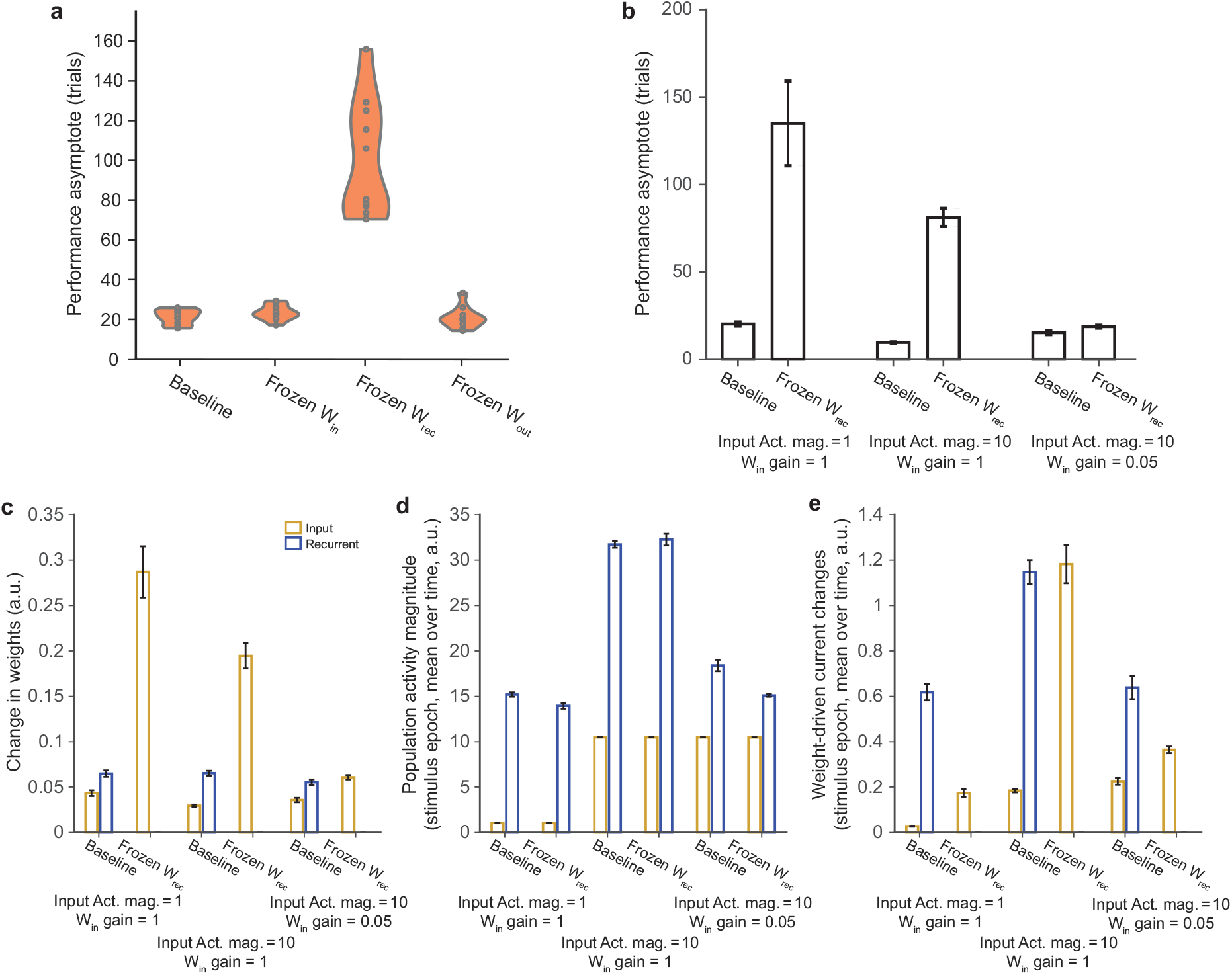
Plasticity of recurrent connection weights results in, but is not necessary for, efficient learning. **a.** Comparison of the learning-to-learn asymptotes of fully plastic networks (Baseline), and those trained with frozen input, recurrent or output weights. Weights were frozen after the first problem was learned, to ensure the emergence of the decision and stimulus manifolds. **b-e.** Comparison of fully plastic networks (baseline) and those with frozen recurrent weights in three network regimes that differed in the magnitude of the input unit firing rates (Input Act. mag.) and the relative strengths of the initial input connection weights (W_in_ gain). Comparison of the learning-to-learn asymptotes (**b**), magnitude (Frobenius norm) of input and recurrent weights changes (**c**), magnitude (L^2^ norm) of input and network unit firing rates (**d**), and magnitude (L^2^ norm) of the input and recurrent postsynaptic current changes (**e**). Quantities in (d-e) are temporal means over the sample epoch of the trials. Quantities in (c-e) are averages across problems 951-1000. All bars represent means across 10 networks with different initial conditions, and error bars indicate their standard errors.

**Supplementary Figure 7:**
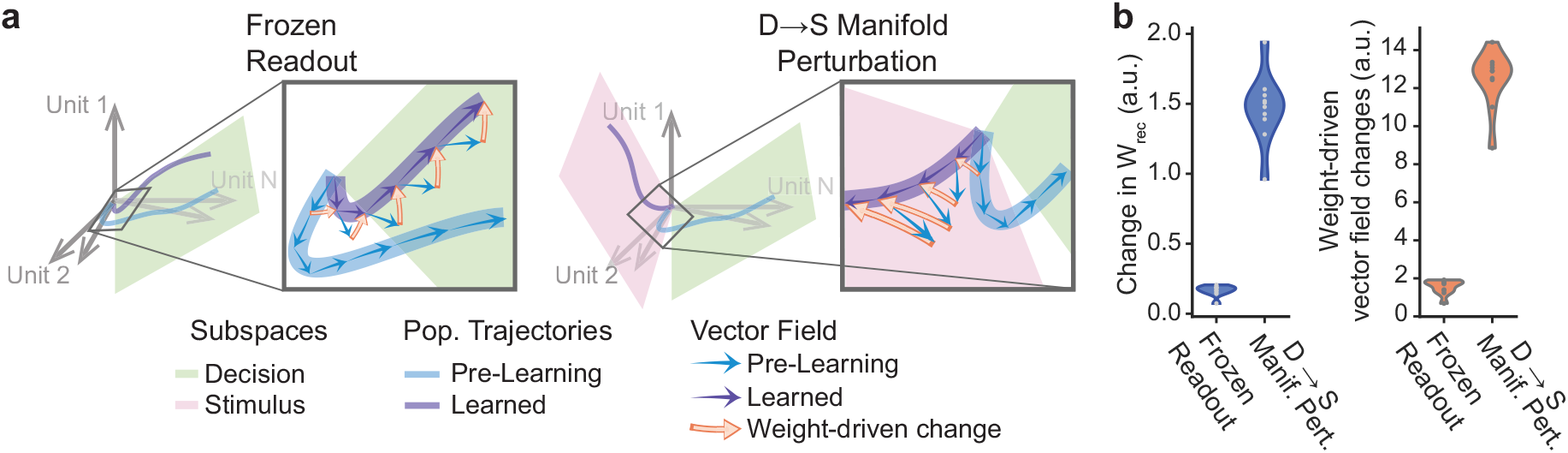
The pre-existing decision manifold provides a representational scaffold for the formation of learned trajectories in new problems. **a.** Schematic illustrating pre-learning and learned population trajectories, together with the requisite weight-driven vector field change (orange arrows) in a network with frozen readouts (left) and a network with a D → S manifold perturbation (right). Vector field shaped by prior learning is approximately oriented to support the evolution of learned trajectories in networks with frozen readouts, but not in networks with D → S manifold perturbations. Therefore, the latter requires a substantial weight-driven vector field change. **b.** Distribution of the magnitude of recurrent weight (left) and weight-driven vector field (right) changes required to learn the second problem, in the frozen readout and D → S manifold perturbation conditions, across 10 networks with different initial conditions. Smaller weight changes required by networks with frozen readouts makes them efficient learners. For each network, the measures were averaged over 50 perturbations, with new sample stimuli used each time. The magnitude (L^2^-norm) of the weight-driven vector field change represented is its temporal mean over the entire trial duration, averaged over both mappings in each problem.

**Supplementary Figure 8:**
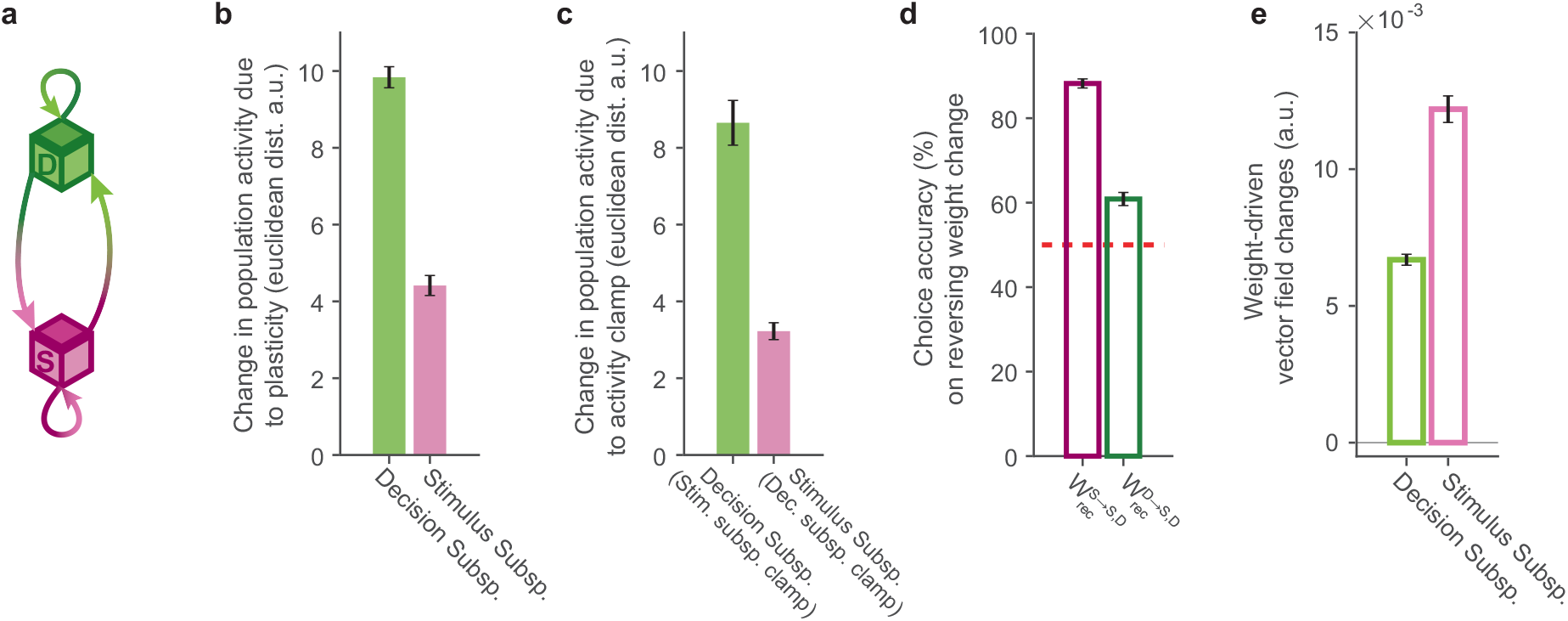
Reciprocal interactions between stimulus and decision representations shape network dynamics and support learning. **a.** Schematic of recurrent interactions within and between stimulus (magenta) and decision (green) representations. Efferent (afferent) connections are represented in darker (lighter) colors. **b.** Euclidean distance between pre-learning and learned decision and stimulus representations. **c.** Euclidean distance between learned decision (stimulus) representations and those generated by the network when simulated with its stimulus (decision) representations clamped to their pre-learning values. **d.** Output accuracy when the weight-driven current changes modulated by presynaptic population activity in the stimulus or decision subspace are reversed. **e.** Magnitude (L^2^-norm) of weight-driven vector field change within the stimulus and decision subspaces. The magnitudes and Euclidean distances represented are their temporal mean over the entire trial duration, averaged over both mappings in problems 2-51. All bars represent mean values over 10 networks with different initial conditions, and error bars indicate their standard errors.

**Supplementary Figure 9:**
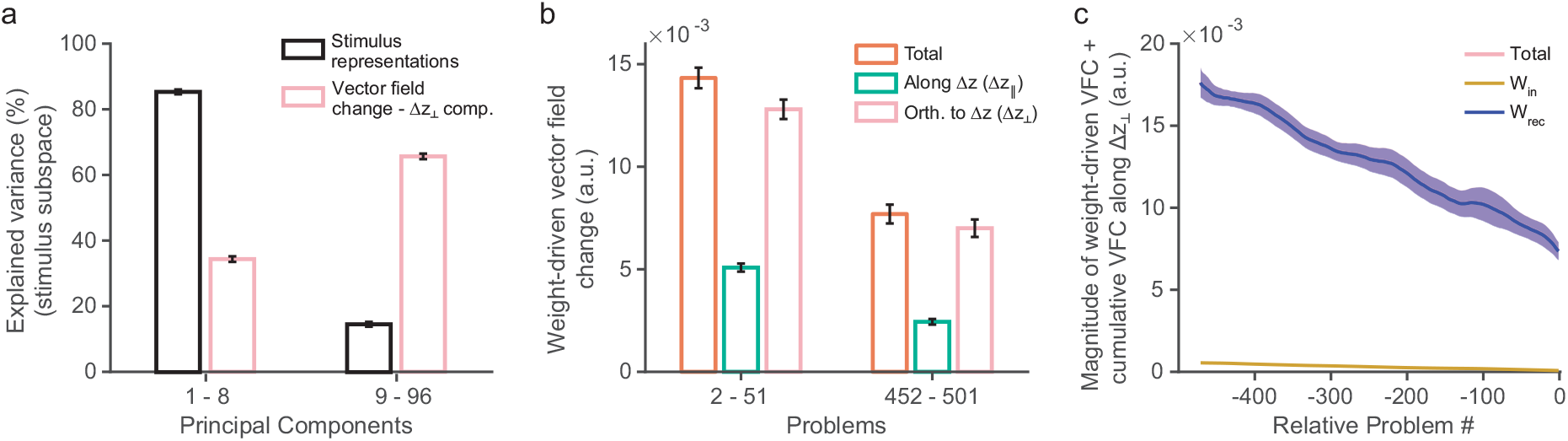
Properties and contributions of the orthogonal components of the vector field change. **a.** Comparison of the percentage of variance in the stimulus representations (black) and the orthogonal components of the vector field change within the stimulus subspace (pink) explained by the first 8 and remaining principal components of the stimulus subspace. The orthogonal components largely lie off the stimulus manifold. **b.** Magnitudes of the weight-driven vector field change (orange), and its components in the direction of the activity change increments (green) and orthogonal them (pink), for early and late learned problem groups. The orthogonal components dominate the total weight-driven vector field change across the learning-to-learn timecourse. **c.** Magnitude of vector field change along the learned trajectory for a problem p due to the accumulation of (i) the weight changes in problem p (W^p^ − W^p−1^, relative problem = 0), and (ii) weight changes in each of the earlier problems, proceeding backwards to problem 2 (W^p^ −W^p−k−1^ for 1 ≤ k ≤ p− 2, relative problem −k). The curve for each problem measures the magnitude of change in the direction of its orthogonal weight-driven VFC component, smoothed with a 30-problem moving average filter. Plot summarizes the measurements for problems at late-stage learning-to-learn (problems 452-501), and separately shows contributions of changes in input weights (yellow), recurrent weights (blue), and both (pink). The cumulative suppression of the weight-driven vector field change in the direction of its orthogonal component is almost entirely caused by an accumulation of recurrent weight changes. The magnitude (L^2^-norm) of the vector field change represented in (b-c) is its temporal mean over the entire trial duration, averaged over both mappings of all the problems in the respective group of 50 problems. All plots represent mean values over 10 networks with different initial conditions, and shading/error bars indicate their standard errors.

**Supplementary Figure 10:**
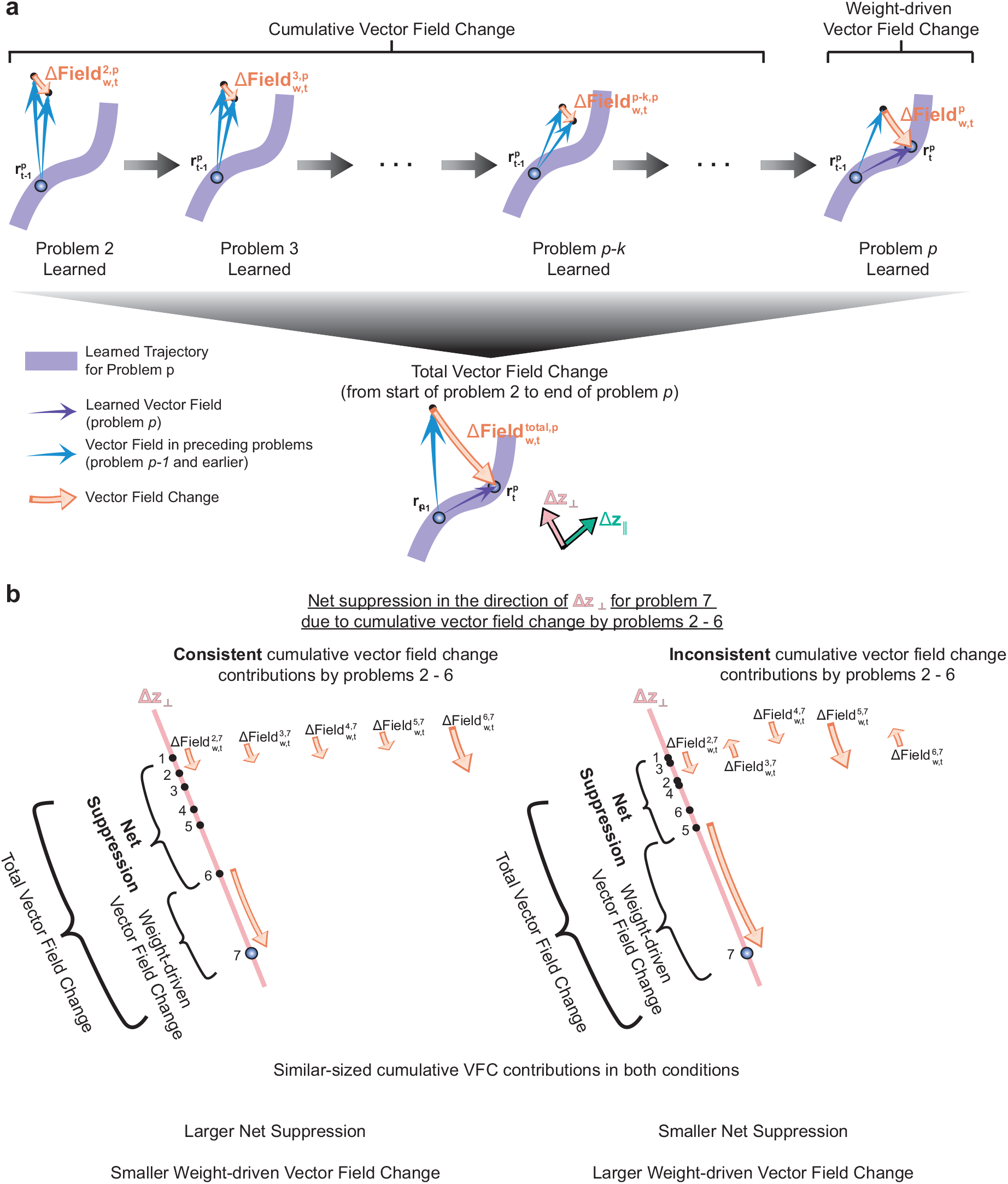
Weight-driven vector field changes in earlier problems cumulatively suppress the weight-driven vector field change required to learn future problems. **a.** Schematic illustrating cumulative changes in the vector field along the learned trajectory for problem p due to an accumulation of the weight changes elicited while learning the problems that precede it. This cumulative vector field change reduces the weight-driven vector field change required to support the evolution of problem p’s learned trajectory. The total vector field change at problem p measures the net effect of all the vector field changes along problem p’s learned trajectory due to the weight changes between the start of problem 2 and the end of problem p. **b.** Difference between consistently and inconsistently suppressive cumulative vector field change contributions illustrated along a problem’s orthogonal vector field change component. Stronger consistency produces a larger net suppression, which reduces the requisite weight-driven vector field change by a larger amount.

**Supplementary Figure 11:**
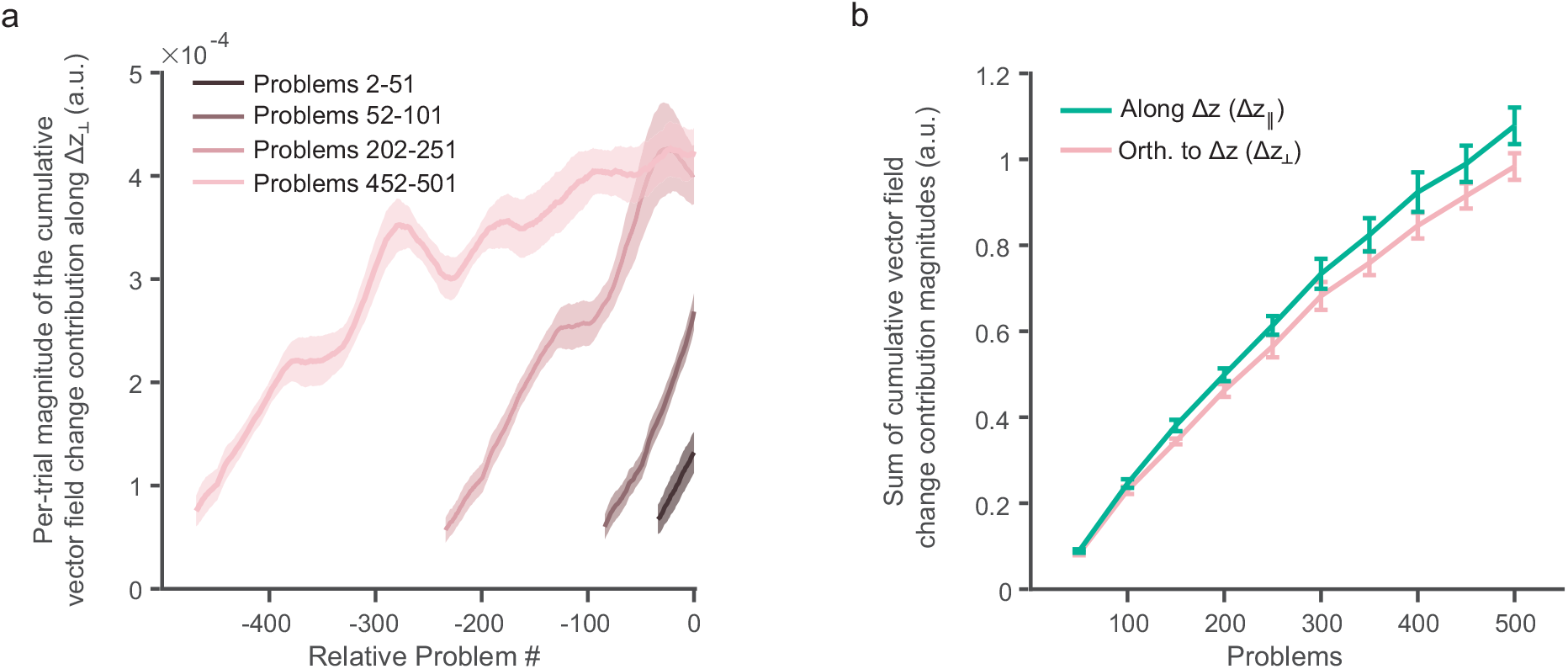
Properties of vector field change along the learned trajectory for a problem due to weight changes in the preceding problems. **a.** Magnitude of per-trial cumulative vector field change contributions along the learned trajectory for a problem p by the weight changes in each of the earlier problems, proceeding backwards to problem 2 (W^p^ − W^p−k−1^ for 1 ≤ k ≤ p − 2, relative problem −k). The curve for each problem measures the per-trial magnitude of change in the direction of its orthogonal weight-driven VFC component, smoothed with a 30-problem moving average filter. Plot summarizes the measurements for problems in 4 problem groups at different stages of learning-to-learn. **b.** Sum of the magnitudes of the cumulative VFC contributions for each problem p due to weight changes in problems 2 thru p − 1. summarized in 50-problem groups. The measure is presented separately for the vector field change along the parallel (green) and orthogonal (pink) weight-driven VFC components. Magnitudes shown in both plots are the temporal mean of the unsigned projections onto the parallel / orthogonal weight-driven VFC components, averaged over both mappings in a problem. Plots reflect mean values over 10 networks with different initial conditions, and shading/error bars indicate standard errors.

